# COMPASS: Component-Wise Inference of Shared and Gene-Specific Perturbation Response

**DOI:** 10.64898/2026.08.03.742643

**Authors:** Huan Liang, Rohit Singh

## Abstract

Predicting how a genetic perturbation reshapes a cell’s transcriptome is a central goal of computational biology. Previous studies report that the mean response across training perturbations rivals specialized models on standard accuracy metrics, even though it cannot distinguish which perturbation occurred. Across 2,270 CRISPRi perturbations measured in each of six cell lines, we show that this apparent paradox reflects a conserved organization of perturbation responses. Perturbations span a continuum from responses strongly aligned with the mean to more targeted responses that depart from it. Crucially, a perturbation’s position along this continuum is conserved across cell lines (Kendall’s *W* = 0.59) and predictable from STRING protein-interaction embeddings (*R*^2^ = 0.35). We formalize this structure with COMPASS, an interpretable linear model that decomposes each response into shared and gene-specific components and estimates them separately. The shared-response component is modeled as a cell-line-wide response scaled by a perturbation-specific coefficient. This coefficient is strongly conserved across cell lines. The residual gene-specific component—which is moderately conserved across cell lines—recovers pathway-level programs. COMPASS outperforms scGPT, CPA, GEARS, GenePert, and State in both response accuracy (de-biased Pearson delta 0.34 vs. ≤ 0.32) and perturbation discrimination (cosine PDS gain 0.23 vs. ≤ 0.08). These results recast perturbation prediction across cellular contexts as component-wise inference, with each component estimated from the evidence best suited to it.

## 1 Introduction

One of the long-standing puzzles in perturbation prediction is the strength of simple baselines. For gene knockdowns measured by Perturb-seq, multiple benchmarking efforts^1, 2^ report that the mean response across training perturbations in the same cell line consistently rivals or outperforms purpose-built models such as GEARS and CPA, single-cell foundation models such as scGPT, and architectures trained across diverse experimental contexts.^3–6^ Systema further showed that much of the apparent performance of these models arises from a systematic expression shift shared across perturbations, rather than from recovery of the response specific to the perturbed gene.^7^

While these observations can be read as a failure of model architectures, they also admit structural explanations. The first is about metrics: the mean baseline is *accurate*, recovering much of the measured perturbation effect, but it is ineffective at *discriminating* one perturbation from another. While early studies prioritized accuracy, discrimination is necessary for a complete assessment. The second is about biology: distinct knockdowns may converge on a common response direction because they perturb overlapping cellular programs. Such a shared response would explain the mean baseline’s high accuracy and provide a scaffold for isolating perturbation-specific effects.

If such shared-response structure exists, individual perturbations should differ systematically in how strongly they engage the shared response. Knockdowns of core growth machinery are natural candidates for broad, stereotyped responses in which large numbers of genes shift together. Perturbations of specific regulatory machinery may instead produce responses that depart from the shared direction. To test this hypothesis, we characterize each perturbation *p* in cell line *c* using its response magnitude *m_cp_* ∈ R and its alignment *a_cp_* ∈ R with the cell line’s mean perturbation direction. We explored how *m* and *a* are distributed and whether this organization is conserved across cell lines.

Across 2,270 perturbations shared by six cell lines from three independent CRISPRi studies,^8–10^ we find that magnitude and alignment trace a similar continuum in every cell line. Strikingly, the same perturbations tend to occupy similar positions across studies, and their representations built from prior knowledge, particularly STRING network embeddings, predict those positions. This conservation motivates the decomposition *z_cp_* = *β_cp_u_c_* + *r_cp_*, where *u_c_* ∈ R*^G^* is the cell line’s shared-response direction, *β_cp_* ∈ R measures how strongly perturbation *p* engages that direction, and *r_cp_* ∈ R*^G^* is the gene-specific residual response. Across cell lines, we compute *β*^-^*_p_* and *g_p_* by averaging *β_cp_* and *r_cp_*, respectively. This converts the empirical observation of a perturbation continuum into two quantities with different conservation properties: *β*^-^*_p_* is strongly conserved and transferable across cell lines, while *g_p_* is less well conserved but still retains meaningful cross-cell structure. Both quantities also carry substantial independent biological information.

The two components can therefore be estimated from different evidence and interpreted independently. This motivates COMPASS, an interpretable linear model that estimates these components under within-cell-line and cross-cell-line regimes and combines them to predict unseen perturbation responses. Across six cell lines, COMPASS substantially outperforms direct perturbation-response models and approaches the accuracy of the strongest fixed-representation predictor, TabICL, while retaining stronger cross-cell discrimination than it.

## 2 Methods and Results

We interleave methods and results because our model is motivated directly by exploratory analysis of perturbation-response patterns.

### 2.1 Data and evaluation contexts

We analyzed six pooled CRISPRi Perturb-seq datasets from three independent studies: K562 and RPE1 from Replogle *et al.*, HepG2 and Jurkat from Nadig *et al.*, and HCT116 and HEK293T from X-Atlas/Orion.^8–10^ After applying each study’s guide-assignment and cell-quality filters, we intersected the targets measured in all six lines to obtain 2,270 shared perturbations, predominantly essential genes from the Cancer Dependency Map project.^11, 12^ The shared perturbation set therefore supports matched cross-cell analysis but is enriched for genes with strong fitness effects, a limitation we return to in the Discussion. Counts were library-size normalized to 10,000 per cell and log-transformed as log(1 + *x*).

Predictions were scored on panels of 2,000 highly variable genes. We used two kinds of panels: cell-line-specific panels, selected independently for each line, and dataset-specific panels, shared across cell lines within a study group. Because the Replogle/Nadig (four cell lines) and Orion (two lines) dataset-specific panels overlap in only 282 readout genes, we kept our analysis dataset-specific; cross-cell quantities were estimated only within these groups. We evaluated predictions in two settings that correspond to different experimental situations. In the *within-cell-line* setting, a model observes a subset of perturbations in the query cell line and predicts held-out perturbations in the same line. This corresponds to extending a partially completed screen. In the *cross-cell-line* setting, the query perturbation has been measured in other cell lines but is held out from the target line. This corresponds to transferring knowledge from existing screens to a new cellular context. Extensive data processing details can be found in Section A.1 and Section A.2.

### 2.2 Evaluation metrics: accuracy and discrimination

A perturbation predictor must do two separable things: recover what the response looks like (*accuracy*) and distinguish one perturbation’s response from another (*discrimination*). Measuring them requires distinct metrics.

For perturbation *p*, cell line *c*, and readout gene *g* ∈ G with |G| = *G*, we define the pseudobulk perturbation effect

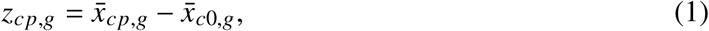

here *x^-^*_c0,*g*_ is the mean among non-targeting controls. A predictive model returns a vector z^*_cp_* ∈ R*^G^* for each held-out perturbation.

We measure accuracy with the **de-biased Pearson delta**, which quantifies agreement across readout genes between the predicted and measured perturbation-effect vectors. Following Nicol *et al.*, we form the predicted and observed effects using disjoint halves of the non-targeting controls, preventing their shared control reference from inflating the correlation.^13^ We measure discrimination with the **cosine perturbation discrimination score (PDS) gain**: PDS tests whether each prediction is closer to its matched measured response than to those of other perturbations. We use cosine similarity, as ℓ_1_/ℓ_2_ is affected by magnitude of predicted vectors.^14^ A score of 0.50 corresponds to chance, and we report the gain above chance.

We also computed DE overlap@*N*, which measures overlap between the top-ranked differential genes in the predicted and measured responses, with *N* set by the number of measured genes passing FDR < 0.05. Because DE overlap ranks methods nearly identically to the de-biased Pearson delta across all methods and cell lines (Table 4), we report it in the supplement as a confirmatory accuracy measure. All formal metric definitions appear in Section A.3.

### 2.3 The mean baseline captures the shared response but cannot discriminate perturbations

The training mean recovers much of the measured response but cannot distinguish which perturbation produced it. Across five random 80/20 splits per cell line, assigning every held-out perturbation the average training response matches or exceeds GEARS and scGPT on de-biased Pearson delta in every cell line, reaching 0.09–0.55 compared with 0.03–0.43 for GEARS (Table 1). By construction, the same prediction is returned for every test perturbation, so its cosine PDS remains at chance (0.50). GEARS and scGPT also remain near chance, at gains of 0.01 and 0.00, despite producing perturbation-conditioned outputs. Their response-level agreement therefore comes primarily from structure shared across perturbations. All models and the training mean are described in Section A.4.

**Table 1:** The mean-baseline puzzle. De-biased Pearson delta and cosine PDS gain above chance on the per-cell 2,000-HVG panels, averaged over five random 80/20 partitions per cell. In each column and metric block, **bold** marks the best value and <u>underline</u> the second best, with all tied values underlined.

|  | K562 | RPE1 | HepG2 | Jurkat | HCT116 | HEK293T | Mean |
| --- | --- | --- | --- | --- | --- | --- | --- |
| **Accuracy: De-biased Pearson delta** |  |  |  |  |  |  |  |
| Training mean | <u>+0.26</u> | <u>+0.55</u> | <u>+0.36</u> | <u>+0.24</u> | <u>+0.19</u> | <u>+0.09</u> | <u>+0.28</u> |
| CPA | +0.05 | +0.06 | +0.06 | +0.07 | +0.01 | +0.02 | +0.05 |
| GEARS | +0.13 | +0.43 | +0.25 | +0.13 | +0.06 | +0.03 | +0.17 |
| scGPT | +0.18 | <u>+0.55</u> | +0.34 | +0.19 | +0.16 | +0.06 | +0.25 |
| GenePert | **+0.32** | **+0.57** | **+0.38** | **+0.28** | **+0.23** | **+0.13** | **+0.32** |
| **Discrimination: Cosine PDS gain** |  |  |  |  |  |  |  |
| Training mean | +0.00 | +0.00 | +0.00 | +0.00 | +0.00 | +0.00 | +0.00 |
| CPA | <u>+0.01</u> | +0.00 | +0.01 | +0.02 | <u>+0.01</u> | +0.00 | <u>+0.01</u> |
| GEARS | +0.00 | <u>+0.01</u> | <u>+0.03</u> | <u>+0.03</u> | -0.01 | <u>+0.02</u> | <u>+0.01</u> |
| scGPT | +0.00 | +0.00 | +0.00 | +0.00 | +0.00 | +0.00 | +0.00 |
| GenePert | **+0.11** | **+0.05** | **+0.06** | **+0.09** | **+0.08** | **+0.07** | **+0.08** |

**Table 2:**
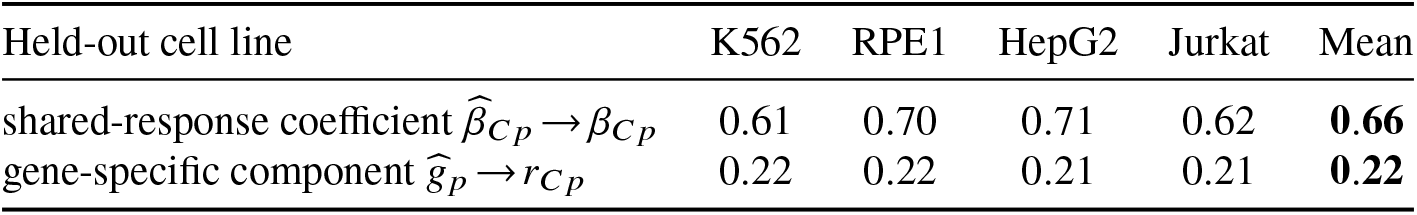
The shared-response coefficient transfers more strongly than the gene-specific response. Each of the four Replogle/Nadig cell lines is held out in turn, and both components are estimated from the other three. The coefficient is a scalar per perturbation, compared by Pearson correlation across the 2,270 perturbations; the gene-specific response is a vector, compared by Pearson correlation across genes within each perturbation and averaged. A perturbation-shuffled null gives ≈ 0 for both.

| Held-out cell line | K562 | RPE1 | HepG2 | Jurkat | Mean |
| --- | --- | --- | --- | --- | --- |
| shared-response coefficient $\widehat{\beta}_{Cp} \rightarrow \beta_{Cp}$ | 0.61 | 0.70 | 0.71 | 0.62 | **0.66** |
| gene-specific component $\widehat{g}_p \rightarrow r_{Cp}$ | 0.22 | 0.22 | 0.21 | 0.21 | **0.22** |

**Table 3:** Cross-context prediction across the six cell lines (random splits). De-biased Pearson delta and cosine PDS gain, averaged over five 80/20 partitions per cell line. Cross-cell methods are scored on the group’s shared panel, within-cell methods on the query cell line’s own panel. TabICL is run with all six prior-knowledge modalities as its authors do. **Bold** marks the best value in each column and metric, <u>underline</u> the second best.

|  |  | K562 | RPE1 | HepG2 | Jurkat | HCT116 | HEK293T | Mean |
| --- | --- | --- | --- | --- | --- | --- | --- | --- |
| **Accuracy: De-biased Pearson delta** |  |  |  |  |  |  |  |  |
| Cross-cell | Source average | 0.28 | 0.37 | 0.35 | 0.26 | 0.10 | 0.08 | 0.24 |
|  | STATE | 0.14 | 0.44 | 0.31 | 0.17 | 0.10 | 0.08 | 0.21 |
|  | TabICL | **0.41** | **0.63** | **0.46** | **0.41** | **0.28** | **0.20** | **0.40** |
|  | COMPASS-X | 0.33 | 0.43 | 0.40 | 0.32 | 0.12 | 0.12 | 0.29 |
|  | COMPASS-H | <u>0.36</u> | <u>0.51</u> | <u>0.43</u> | <u>0.35</u> | <u>0.22</u> | <u>0.16</u> | <u>0.34</u> |
| Within-cell | Training mean | 0.26 | 0.55 | 0.36 | 0.24 | 0.19 | 0.09 | 0.28 |
|  | CPA | 0.05 | 0.06 | 0.06 | 0.07 | 0.01 | 0.02 | 0.05 |
|  | GEARS | 0.13 | 0.43 | 0.25 | 0.13 | 0.06 | 0.03 | 0.17 |
|  | scGPT | 0.18 | 0.55 | 0.34 | 0.19 | 0.16 | 0.06 | 0.25 |
|  | GenePert | 0.32 | 0.57 | 0.38 | 0.28 | 0.23 | 0.13 | 0.32 |
|  | TabICL | **0.39** | **0.61** | **0.43** | **0.35** | **0.28** | **0.19** | **0.37** |
|  | COMPASS-N | <u>0.34</u> | <u>0.57</u> | <u>0.39</u> | <u>0.30</u> | <u>0.25</u> | <u>0.16</u> | <u>0.34</u> |
| **Discrimination: Cosine PDS gain** |  |  |  |  |  |  |  |  |
| Cross-cell | Source average | **0.25** | **0.32** | **0.24** | <u>0.23</u> | <u>0.18</u> | <u>0.16</u> | **0.23** |
|  | STATE | 0.18 | 0.04 | 0.04 | 0.11 | 0.07 | 0.06 | 0.08 |
|  | TabICL | 0.24 | 0.18 | 0.20 | **0.27** | 0.15 | 0.16 | 0.20 |
|  | COMPASS-X | **0.25** | <u>0.30</u> | <u>0.22</u> | 0.24 | <u>0.18</u> | **0.19** | **0.23** |
|  | COMPASS-H | <u>0.24</u> | 0.28 | 0.21 | 0.24 | **0.23** | **0.19** | **0.23** |
| Within-cell | Training mean | 0.00 | 0.00 | 0.00 | 0.00 | 0.00 | 0.00 | 0.00 |
|  | CPA | 0.01 | 0.00 | 0.01 | 0.02 | 0.01 | 0.00 | 0.01 |
|  | GEARS | 0.00 | 0.01 | 0.03 | 0.03 | -0.01 | 0.02 | 0.01 |
|  | scGPT | 0.00 | 0.00 | 0.00 | 0.00 | 0.00 | 0.00 | 0.00 |
|  | GenePert | 0.11 | 0.05 | 0.06 | 0.09 | 0.08 | 0.07 | 0.08 |
|  | TabICL | **0.21** | **0.14** | **0.15** | **0.19** | **0.16** | **0.15** | **0.16** |
|  | COMPASS-N | <u>0.19</u> | <u>0.13</u> | <u>0.13</u> | <u>0.18</u> | <u>0.15</u> | <u>0.12</u> | <u>0.15</u> |

GenePert outperforms the other baselines through a structure consistent with the decomposition we formalize later in this section. The method fits a regularized regression from a gene embedding to the perturbation effect within a cell line and achieves the strongest response correlation among the within-cell baselines (0.32 on average, compared with 0.28 for the training mean). Its fitted intercept is the training mean, so each prediction is the shared response plus a shrunken gene-dependent correction. This gene-dependent correction raises its mean PDS gain to 0.08, although its discrimination remains modest relative to its response correlation.

Together, these results separate two prediction tasks: the mean’s high correlation shows that many perturbations share a dominant expression pattern, while its zero PDS gain shows that the perturbation-specific signal resides in the variation around the shared response. We next characterize each response by both its overall strength and its agreement with the shared direction.

### 2.4 Magnitude and alignment characterize perturbation responses

For each perturbation *p* in cell line *c*, we computed a signed Wilcoxon rank-sum statistic for every readout gene by comparing perturbed cells with non-targeting controls from the same experiment. The resulting profile *w_cp_* ∈ R*^G^* summarizes the direction and consistency of the response. With 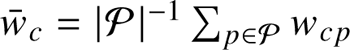 denoting the mean profile over the 2,270 shared perturbations, we define

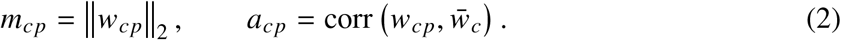

Magnitude measures the overall response strength, while alignment measures how closely the response follows the cell line’s shared perturbation direction.

Magnitude and alignment are positively correlated in all six cell lines (Fig. 2), with Spearman correlations ranging from *ρ* = 0.41 in HEK293T to 0.82 in HepG2. As elaborated in Section 2.6, high-magnitude, high-alignment perturbations elicit broad, similar responses, suggesting extensive disruption of the cellular system. At the other end are perturbations that elicit specific responses that are poorly aligned with the mean.

**Figure 1:**
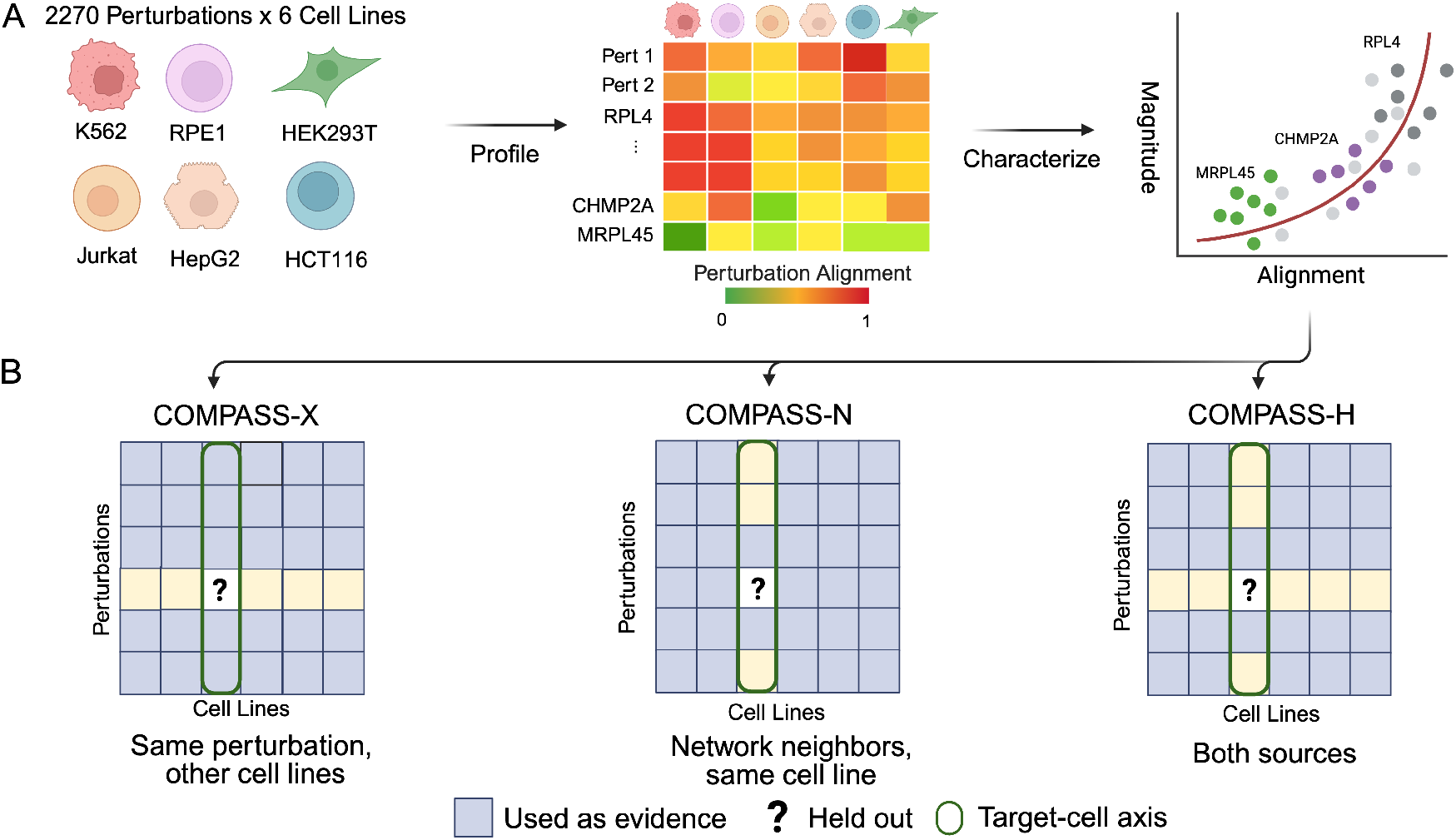
Study overview. **(A)** We profiled 2,270 CRISPRi perturbations shared across six cell lines and characterized each perturbation by its response magnitude and its alignment with the cell-line average response. **(B)** COMPASS predicts unseen perturbations under three evidence regimes: COMPASS-X uses the same perturbation in other cell lines, COMPASS-N uses network-neighbor perturbations in the target cell line, and COMPASS-H combines both.

**Figure 2:**
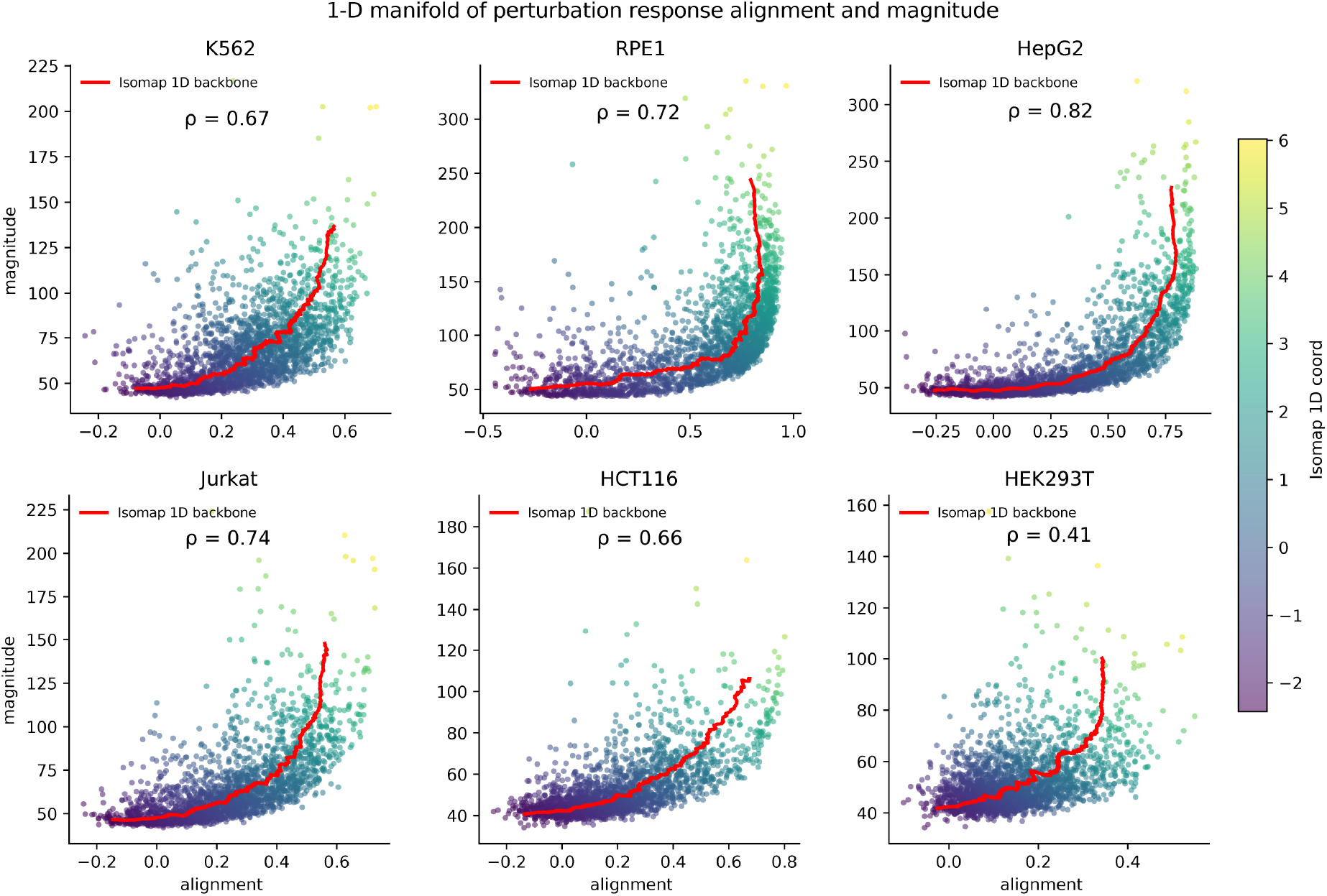
Magnitude and alignment trace a common response organization across six cell lines. Response magnitude *m_cp_* against alignment *a_cp_* for the 2,270 shared perturbations. Cell-specific Spearman correlations between magnitude and alignment are marked on each plot. Points are coloured by the coordinate *s_cp_* of Section 2.5.

This correlation is partly geometric: high-magnitude responses contribute disproportionately to *w^-^_c_* and therefore tend to align with the direction they collectively define, whereas the alignment of weak responses is estimated less reliably. Because magnitude and alignment are naturally coupled, they define a continuum rather than independent axes, motivating a single coordinate along the dominant trajectory (Section 2.5). This structure is robust to the choice of readout panel: recomputing the analysis with 1,000- and 5,000-highly variable gene panels preserves the continuum, cross-cell concordance, and response regimes (Supplementary Fig. 5 and Table 5).

### 2.5 A one-dimensional coordinate orders the magnitude–alignment continuum

To reduce this continuum to a single quantity, we standardized *m_cp_* and *a_cp_* within each cell line and computed a one-dimensional Isomap coordinate, ***s_cp_***, oriented so that larger values of *s_cp_* correspond to the high-magnitude, high-alignment arm (Section A.5). This ordering is conserved across cell lines and studies. Across the fifteen cell-line pairs, Pearson correlations average *r* = 0.50 for alignment, *r* = 0.38 for magnitude, and *r* = 0.50 for *s_cp_*, and every pairwise comparison is positive; Kendall’s coefficients of concordance across all six lines are *W_a_* = 0.587, *W_m_* = 0.533, and *W_s_* = 0.588 for alignment, magnitude, and *s_cp_*, respectively (Fig. 3). Perturbations with high (low) *s_cp_* in one cell line therefore tend to rank highly (lowly) in the others. This conservation persists across study-specific processing pipelines and environmental factors: the strongest concordance in *s_cp_* occurs for the cross-study RPE1–HepG2 pair (*r* = 0.66), and the next strongest, HepG2–HCT116 (*r* = 0.62), is also cross-study, while K562 and RPE1, the two Replogle lines, are among the least concordant (*r* = 0.41, thirteenth of fifteen).

**Figure 3:**
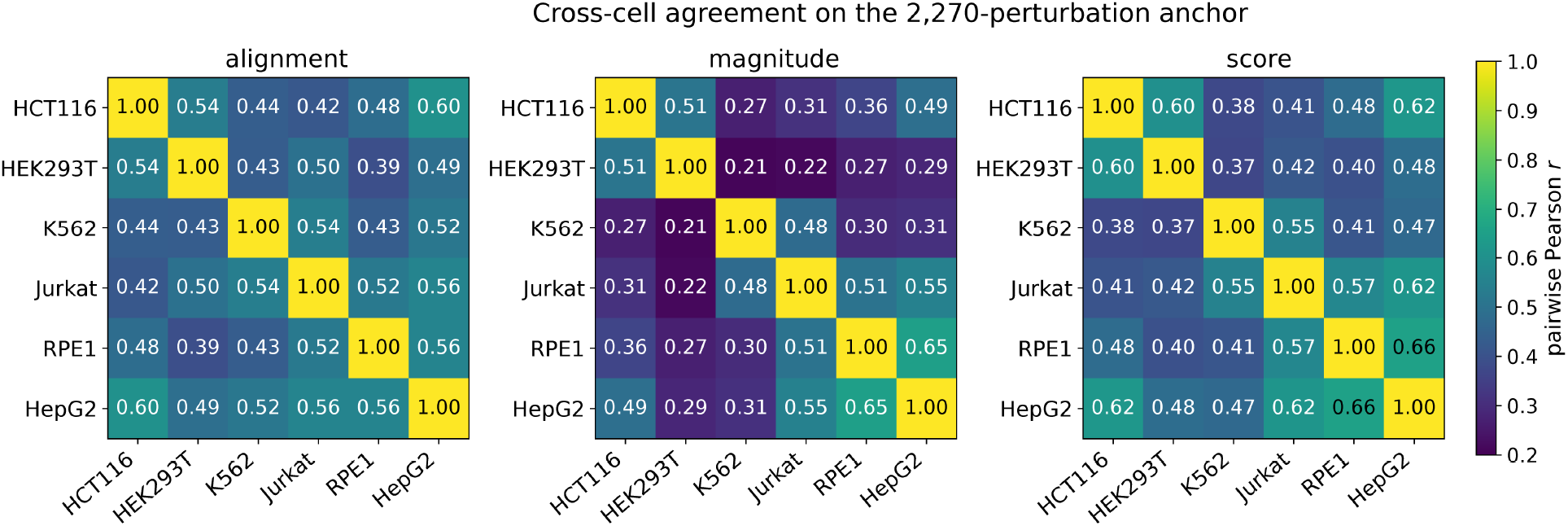
The response organization is conserved across cell lines and studies. Pairwise cross-cell correlations of alignment, magnitude, and the coordinate *s_cp_* across the 2,270 shared perturbations; all fifteen pairs are positive for each measurement.

### 2.6 The two ends of the continuum are biologically distinct

The conserved ordering separates perturbations with distinct biological roles. We averaged the cell-specific coordinates to *s^-^*(*p*) = |C|^−1^ *_c_ s_cp_*, ranked the 2,270 perturbations, and tested the top and bottom 30% for Gene Ontology overrepresentation (Supplementary Table 6). The high-magnitude, high-alignment end is dominated by core growth machinery, including cytoplasmic translation, rRNA processing, spliceosomal mRNA processing, RNA polymerase II preinitiation, and DNA replication. Representative genes include RPL4, RPS13, SF3B3, PRPF31, MCM2, MCM3, MCM6, GINS1, and GINS4. The low-*s^-^* tail is enriched for regulatory and organelle-specific functions, including mitochondrial translation, chromatin remodeling, transcriptional coactivation, protein folding, and regulation of DNA-templated transcription, with genes such as MRPL45, MRPS7, KDM6A, BRD4, NCOA4, and PFDN1. The same terms remain significant when the tails are defined at 20%, 25%, or 30%; the full sensitivity analysis appears in Supplementary Table 7. These enrichments support the biological distinction underlying the continuum: core growth machinery is concentrated among strong, shared-response-dominated perturbations, whereas regulatory and organelle-specific functions are concentrated among less aligned responses.

### 2.7 Gene representations predict conserved response position

Because the same perturbations occupy similar positions across cell lines, we asked whether cell-agnostic prior knowledge about the targeted gene could predict that position. We trained a gradient-boosted regressor to predict the cross-cell means *m^-^* (*p*), *a^-^*(*p*), and *s^-^* (*p*) from gene representations derived from protein-interaction networks, knowledge graphs, transcriptomic data, Gene Ontology annotations, protein sequences, and language models, evaluating every representation under identical perturbation-level five-fold splits, prediction targets, model class, and hyperparameters (Section A.6).

STRING predicts a perturbation’s position more accurately than the other representations (Supplementary Fig. 6). Consistent with its strong cross-cell agreement, alignment is the most predictable of the three quantities, with cross-validated *R*^2^ = 0.39. Magnitude and *s^-^* (*p*) are also well-predicted, albeit less strongly, at *R*^2^ = 0.24 and 0.35, respectively. Graph-neural, knowledge-graph, Gene Ontology, and language-model representations also recover meaningful signal, while somewhat surprisingly, transcriptomic and protein-sequence representations perform less well.

### 2.8 COMPASS: a shared-plus-specific decomposition of the perturbation response

The response structure characterized above motivates a direct decomposition in expression space. For the pseudobulk effect vectors *z_cp_* defined in Eq. (1), 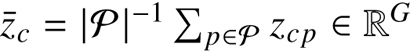 denote the mean perturbation response in cell line *c*, and let 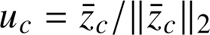 be the unit vector along that mean. We decompose each response as

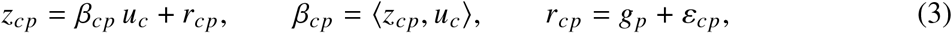

Here *β_cp_* ∈ R is the shared-response coefficient, computed by projecting *z_cp_* onto *u_c_*, and *β_cp_u_c_* ∈ R*^G^* is the corresponding shared-response component. The vector *r_cp_* ∈ R*^G^* is the gene-specific response remaining after this component is removed; it is orthogonal to *u_c_* by construction. Across cell lines with a common readout space, *g_p_* denotes the conserved mean of this gene-specific response and *ε_cp_* its cell-line-specific deviation. A single cell line cannot identify *g_p_* and *ε_cp_* separately, so within-cell estimators target *r_cp_* directly. Here and below, *gene-specific* means specific to the perturbed gene *p*, rather than to an individual readout gene.

The shared-response coefficient *β_cp_* is the expression-space counterpart of the empirical coordinate *s_cp_*: both order perturbations by how strongly their responses follow the cell line’s shared response. *β_cp_* is slightly better conserved across cell lines than *s_cp_* itself: per-cell-line estimated values correlate at mean pairwise Pearson 0.55 among the four Replogle/Nadig lines and 0.56 between the two Orion lines (Supplementary Table 13). The two orderings also agree closely: the Pearson correlation between the (Replogle/Nadig) cross-cell mean *β*^-^*_p_* = |C|^−1^ *β_cp_* and *s^-^* (*p*) is 0.88. Ranking by *β*^-^*_p_* yields similar gene function enrichments (Supplementary Table 8). Because *β*^-^*_p_* can be estimated from source cell lines and summarizes a perturbation’s shared-response engagement in a single portable scalar, it can be used as an input in cross-cell prediction settings.

To measure how well these components transfer across cellular contexts, we performed leave-one-cell-line-out assessments on the four Replogle/Nadig lines, writing *C* for the held-out line. The estimate *β*^^^*_C p_*, formed by averaging over the remaining three lines, predicts the held-out shared-response coefficient *β_Cp_* with mean Pearson correlation 0.66, while the corresponding estimate *g_p_* matches the held-out gene-specific response *r_Cp_* at mean Pearson 0.22 (Table 2). The shared-response coefficient thus transfers considerably more reliably than the gene-specific residual. With only one source line available, as for the Orion pair, transfer remains predictive but is clearly weaker than from three, 0.56 and 0.09 for the two components (Supplementary Table 9).

Together, these analyses identify two components with complementary conservation properties: the shared-response coefficient *β*^-^*_p_* is strongly conserved across cell lines and transfers reliably, while the gene-specific response, though more cell-line-specific, still retains meaningful cross-cell-line structure.

### 2.9 Estimating COMPASS under different information regimes

We now consider predicting *z_Cp_* for an unmeasured perturbation *p* in a target cell line *C*. COMPASS assumes that the target shared-response direction *u_C_* can be estimated from a calibration set of perturbations measured in *C*; the low-data analysis below shows that this estimate stabilizes within a few tens of perturbations (also see Discussion). The prediction task then reduces to estimating *β_Cp_* and *r_Cp_*. COMPASS provides three estimators for these quantities: a cross-cell estimator that carries both components over from other cell lines, a within-cell estimator that infers them from STRING neighbors of *p* measured in *C*, and a hybrid that combines both sources with component-specific weights. In each case, the final prediction is assembled as

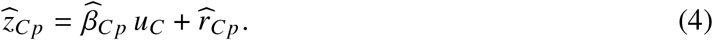

#### COMPASS-X: the perturbation is measured in other cell lines

When *p* is available in a set of source cell lines S*_p_* = {*d* ≠ *C*: *p* ∈ P*_d_* }, we estimate its shared-response projection and conserved gene-specific component by

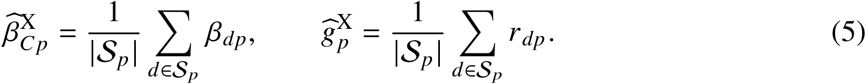

Since the target-specific deviation *ε_Cp_* is unobserved, we approximate 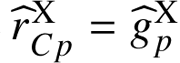 We emphasize that this differs from simply averaging the raw source response vectors *z_dp_* and transferring them directly. COMPASS-X first decomposes each source-line response into its shared-response projection and gene-specific component, then transfers these separately, applying the averaged coefficient 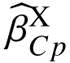 to the target-cell direction *u_C_* rather than retaining the source-cell directions. Critically, this decomposed transfer outperforms direct source averaging, as we show below.

#### COMPASS-N: related perturbations are measured in the target cell

When the target line contains a sufficiently large perturbation set, or when *p* has not been screened elsewhere, we estimate both components from STRING neighbors measured in *C*. With N*_k_* (*p*) denoting the *k* nearest training perturbations,

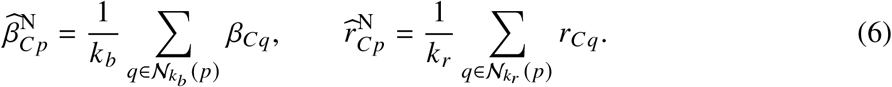

The neighborhood sizes *k_b_* and *k_r_* can be selected separately on validation data (Section A.7); we recommend default values of *k_b_* = 5 and *k_r_* = 10.

#### COMPASS-H: both sources of information are available

When both sources of information are available, we combine the *β* and *r* estimates from COMPASS-X and COMPASS-N. Because the shared-response coefficient and gene-specific response have different transfer properties, COMPASS-H assigns them separate mixing weights:

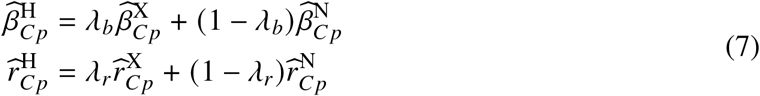

We set *λ_b_* = 1 because the cross-cell estimate comes from the same perturbation in other cell lines and better preserves its magnitude along the shared response direction; a sensitivity analysis confirms that this choice gives the strongest discrimination (Supplementary Fig. 7). We select *λ_r_* jointly with *k_r_* on the validation split, allowing the gene-specific estimate to draw on target-cell neighbors when the conserved component alone is insufficient. As defaults, we recommend *λ_b_* = 1, *λ_r_* = 0.5.

### 2.10 COMPASS offers state-of-the-art perturbation prediction accuracy

To benchmark the predictive performance of our approach, we compare the COMPASS suite against three families of existing models (Table 3): *structural baselines* (training mean, source average), *perturbation-conditioned deep learning models* (CPA, GEARS, scGPT, State), and *embedding-based regressors* (GenePert^15^ and TabICL^16, 17^). Our adaptation of TabICL follows Palla *et al.*’s work. Implementation details for each are given in Section A.4. Compared to these, the COMPASS suite exploits the response structure characterized in Section 2.8: each prediction is a shared-response component plus a gene-specific residual, with both terms estimated from within-cell (COMPASS-N), cross-cell (COMPASS-X) or both (COMPASS-H).

The COMPASS suite, specifically the hybrid model COMPASS-H, matches or outperforms every non-TabICL model on both accuracy (de-biased Pearson delta) and discrimination (cosine PDS gain) (Table 3). COMPASS-X uses the same inputs as the source average but decomposes them first, and is more accurate at equal discrimination. This suggests that transferring information from other cell lines via our linear decomposition is more effective than using it directly. Adding within-cell information, i.e. COMPASS-H, further improves performance, while the source averaging baseline is unable to accommodate such additional information.

COMPASS substantially outperforms the direct perturbation-response deep learning models evaluated here. In within-cell contexts, COMPASS-N leads the deep learning approaches scGPT, GEARS, and CPA. As shown in Section 2.3, several of these models also underperform simple averaging baselines on response accuracy.

GenePert and TabICL are regression-based frameworks that use one and six sets of gene embeddings, respectively, to predict response. GenePert works in a within-cell setting and uses regression; TabICL works in a hybrid setting and uses pre-trained tabular foundation models. We argue they are both best thought of as powerful variants of the training mean or source averaging baseline, as they both learn the overall response and add gene-specific tilts. Interestingly, both methods are stronger on accuracy than discrimination, analogous to the training mean.

TabICL is overall more accurate than COMPASS variants but slightly less discriminative; GenePert is outperformed by COMPASS on both metrics. TabICL leads COMPASS-H on de-biased Pearson delta by 0.06 cross-cell (0.40 vs. 0.34) and COMPASS-N by 0.03 within-cell (0.37 vs. 0.34). On cosine PDS gain, however, COMPASS-X and COMPASS-H match or exceed TabICL cross-cell (0.23 vs. 0.20), and COMPASS-N is within 0.01 within-cell (0.15 vs. 0.16). COMPASS is therefore competitive with a much heavier model, using STRING embeddings only as a distance metric to select neighbors rather than as regression inputs, and its decomposition additionally produces components that are interpretable in isolation (Section 2.11).

#### COMPASS variants reflect the tension between accuracy and discrimination

COMPASS-X is the stronger discriminator (PDS gain 0.23 vs. 0.15) because its perturbation-specific information comes from measurements of the same perturbation in other cell lines. COMPASS-N is the stronger accuracy model (de-biased Pearson 0.34 vs. 0.29) because averaging over STRING neighbors in the target line recovers the local shape of the response, but this neighbor-averaging cannot effectively separate a perturbation from its neighbors. COMPASS-N is therefore the appropriate estimator when the target-cell perturbation set is large but the target perturbation has not been measured elsewhere, while COMPASS-X is the appropriate estimator when the same perturbation has been measured in a related cell line but the target perturbation set is sparse (Supplementary Fig. 8). COMPASS-H estimates each component from the source that recovers it better, matching the stronger of the two on both axes at once.

#### For a new cell line, COMPASS is more data-efficient than TabICL

When investigating a new cell line or tissue, a practical question is how many perturbations are needed before data from other cell lines can be effectively used. To assess this, we included all available cross-cell perturbations and increased the number of within-cell training perturbations from 10 to 130. Across ten seeds in each of the six cell lines (Fig. 4), COMPASS-X converges quickly: its de-biased Pearson delta rises from 0.253 to 0.288 across the range and its PDS gain stays flat near 0.24. This is because the target cell only enters through the shared response *u_C_*, which converges within a few tens of perturbations. In contrast, TabICL starts well below COMPASS-X, and only overtakes it on accuracy at approximately 80 profiled perturbations; it does not outperform on discrimination even with maximal data in this scenario.

**Figure 4:**
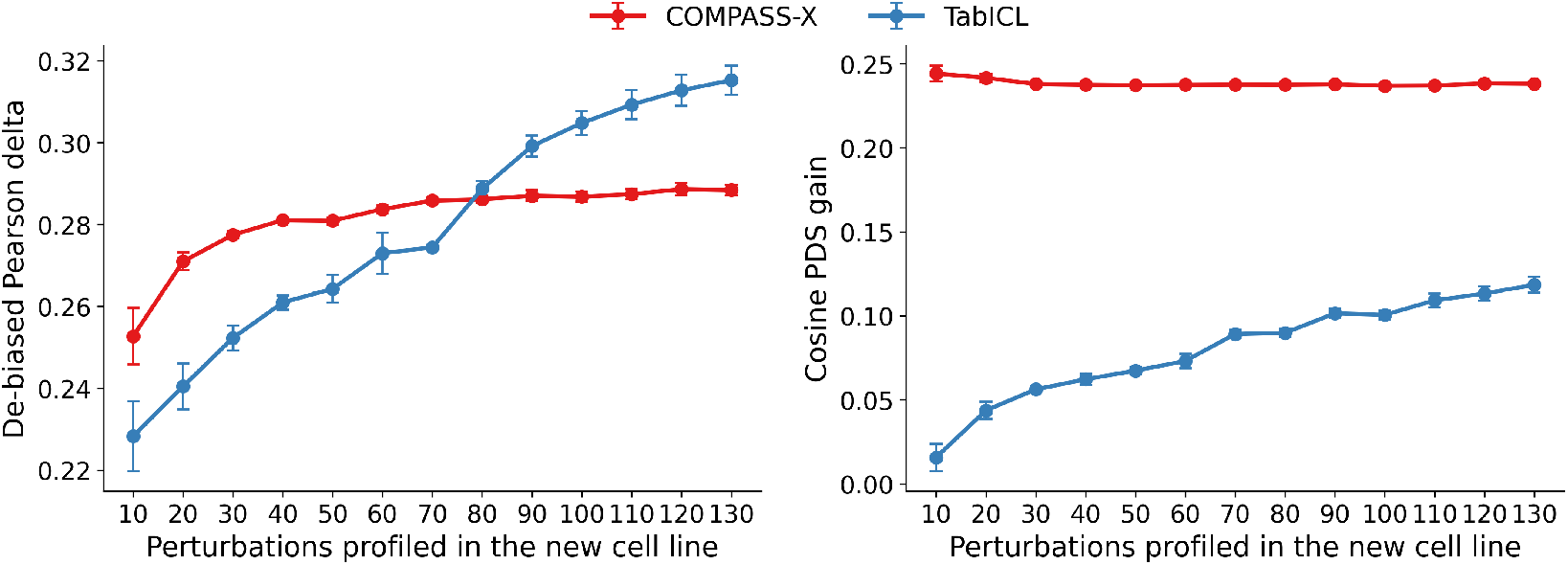
Cross-context transfer versus in-context learning as the profiling budget grows. De-biased Pearson delta (left) and cosine PDS gain (right) against the number of perturbations profiled in a new cell line, with the remainder held out for test.

### 2.11 The gene-specific response ***g_p_*** encodes pathway-level biology

The gene-specific component contains coherent biological programs rather than unstructured variation left after removing the shared response (Section A.8). Averaging *r_cp_* across the four Replogle/Nadig cell lines gives a 2,270 × 2,000 matrix of conserved components *g_p_*. Clustering their normalized directions groups perturbations targeting common biological machinery, including mitochondrial translation, ribosome biogenesis, rRNA processing, mitosis, DNA replication, RNA splicing, and proteasome function (Supplementary Table 10). All fifteen clusters in the highest-coefficient tertile are enriched for perturbations targeting a coherent pathway or complex (Supplementary Table 12). The organization persists among perturbations that engage the shared response weakly, where clusters recover mitochondrial translation, N-glycosylation, and mitochondrial organization programs (Supplementary Table 11).

Individual gene-specific components also recover expected downstream responses. The TFAM component is dominated by reduced expression of mitochondrially encoded oxidative-phosphorylation transcripts. Proteasome-subunit perturbations, including PSMB5, repress cytoplasmic translation while inducing vacuolar and protein-stress programs; the PSMB5 component alone is enriched for unfolded-protein (*q* = 8 × 10^−5^) and heat-shock (*q* = 3 × 10^−5^) responses among its fifteen most induced genes. None of these analyses uses pathway labels during fitting. The gene-specific component therefore captures both relationships among perturbed genes and the downstream programs that distinguish their responses.

## 3 Discussion

The key conceptual advancement of our work is to efficiently disentangle gene-specific effects from shared responses in single-cell perturbation data. This separation both clarifies which pathways a perturbation engages and improves prediction of unmeasured perturbations. Across 2,270 CRISPRi knockdowns, response magnitude and alignment define a continuum from shared-response-dominated perturbations to those eliciting more gene-specific responses. The ordering of perturbations in the continuum is conserved across cell lines and studies. COMPASS formalizes this organization as *z_cp_* = *β_cp_u_c_* + *r_cp_*, with *r_cp_* = *g_p_* + *ε_cp_* across cell lines sharing a readout space. Averaging the shared-response coefficient for the same perturbation across source cell lines yields the portable perturbation-level quantity *β*^-^*_p_*, which predicts how strongly that perturbation will engage the shared response in a new cellular context. The shared-response coefficient transfers substantially more strongly than the gene-specific response *g_p_*, which retains coherent pathway-level structure. Concurrent work by Molina and Zhang^18^ similarly decomposes shared and perturbation-specific response structure, but our work additionally shows that the perturbation-specific coefficient *β_cp_* on the shared response is itself portable across cell lines, enabling COMPASS to reconstruct the shared response as *β*^-^*_p_ u_c_*, contributing to its efficiency and interpretability.

The decomposition also explains why accuracy and discrimination diverge in recent perturbation-prediction benchmarks.^1, 2, 7^ Accuracy metrics reward recovery of the response shared across many perturbations, whereas discrimination rewards capturing what makes each perturbation’s response distinct. The COMPASS variants expose a corresponding tension between sources of evidence. In their corresponding regimes, COMPASS-X and COMPASS-N outperform scGPT, CPA, GEARS, GenePert, and State on both metrics. COMPASS-H combines the strengths of COMPASS-X and COMPASS-N and approaches TabICL’s higher accuracy, exceeds its cross-cell discrimination, and is also more data and parameter-efficient than TabICL.

Our work suggests several directions for future work. As others have noted, perturbation predictors should be evaluated along two distinct axes, response accuracy and perturbation discrimination. Our results further suggest stratifying both metrics along the magnitude–alignment continuum, and model development should similarly treat the two components separately: cross-cell measurements and cell-agnostic gene knowledge may be particularly effective for estimating *β*^-^*_p_*, whereas target-cell data may be needed to refine *r_cp_*. The continuum also suggests a strategy for experimental design. A relatively small set of high *β_p_* perturbations can establish the shared-response direction in a new cell type, allowing more of the profiling budget to be directed toward low-alignment perturbations whose responses carry distinctive and less predictable biology.

There are some limitations to this work. The current perturbation set is enriched for essential genes, so testing broader and more functionally diverse perturbations will establish whether the same shared-plus-specific organization persists beyond stress-dominated responses. Extending the analysis to primary cells and additional cellular contexts will similarly test how well *β*^-^*_p_* and *g_p_* transfer beyond immortalized and cancer-derived cell lines. Finally, COMPASS currently requires perturbations in the target cell line to estimate *u_c_*; learning this direction from unperturbed cells, for example through single-cell foundation-model embeddings, could enable prediction in entirely new cellular contexts without a perturbation calibration set.

## Code and data availability

Code, processed response matrices, split definitions, and scripts for reproducing the figures and tables will be released at https://github.com/rohitsinghlab/compass. Source Perturb-seq datasets remain available from their respective studies.

## Acknowledgements

The authors thank the Whitehead Fund at Duke University for support.

## Author contributions

Both authors conceived of the project. H.L. implemented the software and performed the experiments. Both authors analyzed results and wrote the manuscript.

## A Supplementary Methods

### A.1 Datasets and shared perturbation anchor

We analyzed pooled CRISPRi Perturb-seq screens from K562 and RPE1,^8^ HepG2 and Jurkat,^9^ and HCT116 and HEK293T.^10^ Each dataset contains single-cell expression profiles, guide identities, and non-targeting controls. We restricted every dataset to protein-coding genes using a local copy of the HGNC gene table. For the Ensembl-keyed datasets we stripped the version suffix from each identifier, mapped it to an HGNC symbol, and dropped genes with no mapping. Cells left with zero total counts after this filter were removed.

Control cells were those labelled non-targeting in the source annotation. In K562, HCT116, and HEK293T we reduced the control pool to 10,000 cells before testing; RPE1, HepG2, and Jurkat kept all of their controls (11,485, 4,976, and 12,013 cells). Controls were subsampled by running PCA with 20 components on the normalized control cells, clustering those components with *k*-means into 30 clusters, drawing from each cluster in proportion to its size, and then trimming or topping up at random to reach exactly 10,000 cells. To check this choice we repeated the subsample with five seeds and measured the standard error of each gene’s mean control expression across seeds. We compared stratified against uniformly random sampling of the same pool, and scanned 10, 20, and 50 PCA components against 15, 30, and 60 clusters. The median per-gene standard error under stratified sampling was 0.98–0.99 times the random value in all three cell lines, and varied by at most 3% across the nine grid points.

We retained perturbations assigned at least two cells. Intersecting the retained targets across all six cell lines gave the 2,270-gene shared anchor used for cross-cell analyses.

### A.2 Expression preprocessing and perturbation-effect vectors

Within each cell line, counts were library-size normalized to 10,000 counts per cell and transformed as log(1 + *x*). Highly variable genes were selected with the Seurat normalized-dispersion criterion, taking the top 5,000 genes and recording their rank. We computed two such rankings: one within each cell line, and one across the cell lines of each study group. Analyses use the top 2,000 genes of the relevant ranking.

For response-geometry analyses we compared the cells assigned each perturbation against the control pool with a Mann–Whitney rank-sum test, applying the tie correction and taking the control pool as the reference group. This produced one *z* statistic per gene, giving a response matrix *W_c_* ∈ R*^P^*^×*G*^*^c^* per cell line. The pseudobulk perturbation effect *z_cp_* was computed as in Eq. (1). For benchmark models we used 2,000 highly variable genes and appended perturbed-gene readouts when required by a model implementation. Cross-cell factor models were fit only on common readout panels: a shared 2,000-gene panel for K562, RPE1, HepG2, and Jurkat and a separate common 2,000-gene panel for HCT116 and HEK293T.

### A.3 Evaluation metric definitions

All three metrics compare a predicted perturbation effect with the measured one on the readout panel G, and are computed per perturbation and then averaged over the test set P_test_, with *P* = |P_test_|. Writing *x^-^_cp_* for the mean profile of the cells assigned perturbation *p* in cell line *c*, *x^-^_cp_* for the model’s prediction of that profile, and *x^-^_c_*_0_ for the mean profile of the control cells, the measured effect is *z_cp_* = *x^-^_cp_* − *x^-^_c_*_0_ as in Eq. (1) and the predicted effect is ^*z_cp_* = ^^^*x^-^_cp_* − *x^-^_c_*_0_.

#### De-biased Pearson delta

Referencing both effects to the same *x^-^_c_*_0_ makes the two vectors share a term, which correlates them even when the prediction carries no information.^13^ We therefore split the control cells of cell line *c* into two disjoint halves and use their means 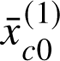 and 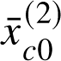 on opposite sides of the comparison,

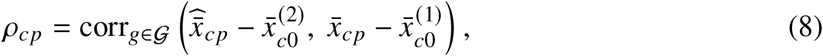

where the correlation runs over readout genes. We draw 20 random halvings of the control pool, use the same 20 halvings for every perturbation, and report the mean of *ρ_cp_* over them.

#### Cosine PDS gain

For each prediction we calculate its cosine similarity to every measured effect in the test pool, and record the rank of its own true measured effect among them. With *k_cp_* being the number of perturbations whose measured effect is closer to the prediction than the matched one is,

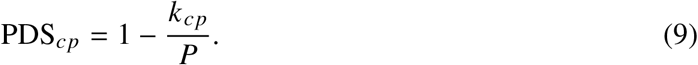

The perturbed gene itself is dropped from G before the distances are computed. A prediction that ranks the matched effect first scores 1, and a random ordering scores 0.5 in expectation; we report the gain PDS*_cp_* − 0.5, averaged across all test perturbations. We directly use the virtual cell challenge’s implementation of this metric.^6^

#### DE overlap@*N*

Differential expression is computed separately for the measured and the predicted responses with the Wilcoxon rank-sum test. Within each, genes are restricted to those passing a Benjamini–Hochberg FDR of 0.05 and ranked by absolute log fold change. We let *N_cp_* to be the **number of measured genes** passing that threshold, and *R_cp_* and *R_cp_* for the actual top *N_cp_* **genes** of the measured and predicted rankings,

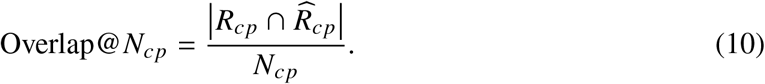

We directly use the virtual cell challenge’s implementation of this metric.^6^

### A.4 Training mean and benchmark predictors

For a training perturbation set P_train_, the training mean predicts every held-out perturbation with

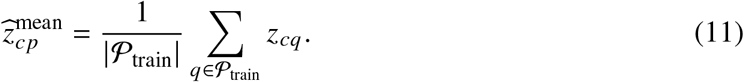

Every other model was run per (cell line, split), trained only on that split’s training perturbations, and asked to predict the same held-out perturbations. We used the scPerturBench implementations and their published hyperparameters,^2^ following each model’s own tutorial and reference script as closely as we could.

#### Source average

The source average is the cross-cell counterpart of the training mean, and like it has no learned parameters and no pretraining. For a held-out cell line it takes the measured effect of the same perturbation in the other cell lines of the same study, averages those effect vectors, and returns the average as the prediction, without separating the shared and gene-specific components. Held-out lines from the Replogle/Nadig group therefore draw on three source lines and the Orion lines on one, and averaging is done on the panel shared within a study group so that every gene is covered in every source. It is the natural control for COMPASS-X, which is given the same information and differs only in decomposing each source response before transferring it.

#### GEARS

GEARS, from Roohani, Huang and Leskovec,^4^ predicts the post-perturbation profile with a graph neural network. Each gene is a node in two graphs, one built from Gene Ontology relations between perturbed genes and one from expression correlation between readout genes, and the network passes messages over both before decoding a per-gene response. It is not pretrained: a fresh model is trained for every cell line and split. We trained with the authors’ settings, Adam at learning rate 10^−3^ with weight decay 5 × 10^−4^, batch size 32, and a step schedule halving the learning rate each epoch, and kept the checkpoint with the lowest validation error on the top 20 differentially expressed genes. We trained for 5 epochs, chosen because that validation error was already lowest at epoch 1 to 3 in every run we inspected and fell by about 1% between epochs 5 and 10. Inference runs the trained network once per held-out perturbation.

#### scGPT

scGPT,^5^ is a transformer trained on tens of millions of single-cell profiles, which is then fine-tuned for a specific task. We started from the authors’ released pretrained weights and fine-tuned separately for each cell line and split. Each training example is a cell, tokenized as a set of genes with their expression values; we used a sequence length of 1,536, sampling genes at random per batch but always including the perturbed gene, with batch size 64 for 10 epochs. The checkpoint was selected on validation correlation of the predicted change. Inference applies the fine-tuned model to the control cells of the target cell line under the held-out perturbation label.

#### GenePert

GenePert,^15^ is a linear model rather than a network: it fits a ridge regression from a fixed embedding of the perturbed gene to that perturbation’s mean expression profile. There is no pretraining step of our own, since the embeddings are precomputed text embeddings of each gene. We fit ridge regression with a penalty of 1 on the training perturbations of each split, using each perturbation’s mean profile as the target. Inference evaluates the fitted regression on the embedding of the held-out gene and returns that profile.

#### CPA

CPA,^3^ is an autoencoder that separates a cell’s expression into a basal state and additive perturbation and covariate effects, with an adversarial term that discourages perturbation information from remaining in the basal representation. We ran the released implementation, including the frozen scGPT gene embedding it ships with, training one model per cell line and split. The reference settings pre-train the autoencoder for thirty epochs, check the validation objective every five epochs, and stop after five checks without improvement. That objective was highest at the first check in every one of our runs, so the restored checkpoint is an early-stopped autoencoder and the adversarial term did not influence any delivered model.

#### State

State,^6^ predicts how a set of cells shifts under a perturbation, and is designed to transfer across cell contexts. We used it as the cross-cell baseline. For each held-out cell line we trained on all anchor perturbations in the other cell lines of the same study group, plus the held-out cell line’s own training perturbations, and predicted its held-out perturbations. Training stayed within a study group so that every cell line shares one gene panel, the group’s top 2,000 highly variable genes, supplied in our normalized space rather than the model’s own. We used the authors’ default optimization settings throughout. Inference predicts the held-out perturbations in the target cell line.

#### TabICL

TabICL, from Qu and colleagues,^16, 17^ is a tabular in-context regressor. Its weights are pretrained on synthetic tables and we applied them frozen, taking no gradient step on our data. Prediction is in-context rather than zero-shot: the training perturbations of a fold are supplied as labelled example rows and the model regresses over them in a single forward pass, so it uses our data every run but retains nothing between folds. Each perturbation is described by six published gene-feature sets covering imaging morphology, language-model, protein-sequence, protein-interaction, and gene-dependency features.^17^ The model predicts in the space of the first 128 principal components of the training-fold responses, and the prediction is mapped back to the readout panel. We ran two settings: a within-cell setting using only these features, and a cross-cell setting that additionally supplies the measured effect of the same perturbation in the other cell lines, with the target cell excluded. A third arm restricted the features to the two protein-interaction sets alone.

### A.5 Magnitude, alignment, and the one-dimensional Isomap coordinate

For each perturbation, magnitude and alignment were computed as in Eq. (2). Within each cell line we took the pair (*a_cp_*, *m_cp_*) over the shared anchor, dropped perturbations missing either value, and standardized each of the two columns to zero mean and unit variance. We then fit a one-dimensional Isomap of the standardized pairs with a neighborhood size of 30. The 1D Isomap is defined only up to a reflection, so we fixed its direction to increase with magnitude, putting the high-magnitude, high-alignment arm at the top of the scale. The cross-cell coordinate is the mean of the six per-cell coordinates.

### A.6 Gene embeddings predict cross-cell response geometry

We asked how well prior knowledge about a gene predicts where its perturbation sits on the response geometry. The gene representations were taken from the collection assembled by Cole *et al.*,^19^ which gathers published embeddings on a common gene index; the STRING protein-interaction embeddings derive from.^20^ We evaluated 57 embeddings in total, spanning protein-interaction networks, knowledge graphs, graph neural networks, Gene Ontology annotations, language models applied to gene descriptions, protein sequence and structure, DNA sequence, and models trained on single-cell expression. Of these, 32 are built in part from cell-line-specific measurements such as essentiality screens. Because those can carry information about the same cell lines we are predicting, we separate them from the 25 cell-agnostic embeddings and report the cell-agnostic ones as the headline comparison.

The three prediction targets were the cross-cell means of magnitude, alignment, and the one-dimensional Isomap. Every embedding was used at its native dimension with no reduction, and every embedding was scored with the same regressor and the same folds: gradient-boosted trees with 300 estimators, maximum depth 6, and learning rate 0.05, under five-fold cross-validation with shuffling and a fixed seed. The folds are over perturbations, so no gene appears in both the training and test halves of a split. Embeddings do not all cover the full anchor, ranging from 2,149 to 2,257 of the 2,270 perturbations, and each was scored on the perturbations it covers. We report the cross-validated *R*^2^.

### A.7 STRING-neighbor estimator

For each test perturbation *p*, we identified its nearest training perturbations by cosine distance in the 512-dimensional STRING embedding. COMPASS-N estimates *β_Cp_* from the mean coefficient of the *k_b_* nearest neighbors and *r_Cp_* from the mean target-cell residual of the *k_r_* nearest neighbors, as in Eq. (6). The two neighborhood sizes were selected jointly on the validation partition, by grid search over *k_b_*, *k_r_* ∈ {3, 5, 10, 20, 50, 100}. The held-out perturbation was excluded from all fitted quantities.

### A.8 Clustering the gene-specific component

For the analyses of Section 2.11, we averaged *r_cp_* over the four Replogle/Nadig cell lines, which share a readout panel, giving one 2,270 × 2,000 matrix of conserved components *g_p_*. Rows were scaled to unit length, so perturbations are compared by which genes they move rather than by how strongly, and clustered with *k*-means: all 2,270 perturbations into 20 clusters, and the top and bottom tertiles of the shared-response coefficient, 749 perturbations each, into 15. The cluster counts were fixed in advance.

Each cluster was characterized on two sides: its member genes against the 2,270-perturbation anchor, and the fifteen most up- and down-regulated genes of its mean component against the 2,000-gene readout panel. Both use a hypergeometric test with Benjamini–Hochberg correction against the Enrichr GO Biological Process 2021, KEGG 2021, Reactome 2022, and MSigDB Hallmark 2020 libraries. The tables report one representative significant term per cell, chosen for interpretability where the top-ranked term was an artefact of gene-set overlap, and left blank where no term reaches *q* < 10^−3^. Single perturbations were tested the same way, on the most changed genes of their own component; those enrichments depend on how many genes are taken, so we report them as directions of effect rather than exact significance levels.

### A.9 Statistical analysis

Cross-cell agreement was summarized by pairwise Pearson correlation and Kendall’s coefficient of concordance *W*; within-cell relationships between magnitude and alignment were summarized by Spearman correlation. Gene Ontology enrichment used a hypergeometric test with Benjamini– Hochberg control of the false-discovery rate.

Benchmark results are averaged over five random 80/20 partitions per cell line, and the low-data sweeps were made over ten draws per setting. Uncertainty is taken across independent splits and cell lines, never across the genes within a single response vector, since those are not independent. Where error bars appear on a curve, each cell line’s own mean across the swept variable is removed before the standard error is taken across cell lines, so the bar reflects how consistently the shape of the curve holds between cell lines rather than how far apart the cell lines sit. In the benchmark tables we report point estimates rather than significance tests for method comparisons, marking the best value in each column and block in bold and the second best underlined, with ties shared.

## B Supplementary tables and figures

### B.1 Differential-expression recovery on random splits

**Table 4:** DE overlap@. *N* **on random splits**, companion to Table 3. Averaged over five 80/20 partitions per cell line, with *N* set per perturbation by the number of measured genes passing FDR < 0.05. Cross-cell methods are scored on the group’s shared highly variable genes panel, within-cell methods on the query cell line’s own panel. **Bold** marks the best value in each column and block, <u>underline</u> the second best.

|  |  | K562 | RPE1 | HepG2 | Jurkat | HCT116 | HEK293T | Mean |
| --- | --- | --- | --- | --- | --- | --- | --- | --- |
| **DE overlap@N** |  |  |  |  |  |  |  |  |
| Cross-cell | Source average | 0.10 | 0.16 | 0.13 | 0.09 | **0.02** | **0.05** | 0.09 |
|  | STATE | 0.04 | 0.14 | 0.10 | 0.04 | 0.01 | 0.01 | 0.06 |
|  | TabICL | **0.18** | **0.33** | **0.19** | **0.16** | 0.01 | 0.02 | **0.15** |
|  | COMPASS-X | 0.12 | 0.17 | 0.15 | 0.10 | **0.02** | <u>0.03</u> | 0.10 |
|  | COMPASS-H | <u>0.14</u> | <u>0.22</u> | <u>0.17</u> | <u>0.11</u> | **0.02** | <u>0.03</u> | <u>0.11</u> |
| Within-cell | Training mean | 0.10 | 0.27 | 0.14 | 0.07 | 0.01 | 0.01 | 0.10 |
|  | CPA | 0.06 | 0.17 | 0.10 | 0.04 | 0.01 | 0.01 | 0.06 |
|  | GEARS | 0.05 | 0.19 | 0.11 | 0.03 | 0.01 | 0.00 | 0.06 |
|  | scGPT | 0.10 | 0.29 | 0.13 | 0.06 | 0.00 | 0.01 | 0.10 |
|  | GenePert | 0.12 | 0.29 | 0.16 | 0.08 | 0.01 | 0.01 | 0.11 |
|  | TabICL | **0.16** | **0.33** | **0.18** | **0.12** | 0.01 | 0.01 | **0.14** |
|  | COMPASS-N | <u>0.14</u> | <u>0.30</u> | <u>0.17</u> | <u>0.10</u> | **0.02** | **0.02** | <u>0.12</u> |

### B.2 The response geometry is robust to the size of the readout panel

All main-text analyses use a 2,000-gene highly variable panel. Because the 1,000-, 2,000-, and 5,000-gene panels are strictly nested within the union set used to compute the per-perturbation rank-sum statistics, we recomputed the geometry at each size without repeating the differential tests (Table 5). Magnitude, alignment, and the derived coordinate *s_cp_* agree closely across panel sizes in every cell. Cross-cell concordance of *s_c_* remains nearly constant (Kendall’s *W* = 0.58–0.59). Alignment concordance decreases slightly as the panel grows, while magnitude concordance increases, making the combined coordinate the most stable summary. The organization is therefore not specific to the 2,000-gene choice.Gene Ontology enrichment at the two ends and its robustness to the tail cutoff

**Table 5:** Panel-size sensitivity of the response geometry. Coordinates were recomputed at 1,000-, 2,000-, and 5,000-gene per-cell highly variable panels from the same per-perturbation rank-sum statistics, over the 2,270 shared perturbations. (a) Within-cell agreement between panel sizes: Pearson *r* for alignment *a* and magnitude *m*, Spearman *ρ* for the derived coordinate *s*. (b) Kendall’s coefficient of concordance across the six cell lines, computed separately at each panel size; the 2,000-gene column reproduces the values reported in the main text. HEK293T is the least stable cell in alignment.

| (a) Agreement between panel sizes, within each cell |  |  |  |  |  |  |  |  |  |
| --- | --- | --- | --- | --- | --- | --- | --- | --- | --- |
| Cell | 1,000 vs 2,000 |  |  | 2,000 vs 5,000 |  |  | 1,000 vs 5,000 |  |  |
| | $a$ | $m$ | $s$ | $a$ | $m$ | $s$ | $a$ | $m$ | $s$ |
| K562 | 0.992 | 0.996 | 0.997 | 0.984 | 0.990 | 0.995 | 0.959 | 0.976 | 0.989 |
| RPE1 | 0.998 | 0.998 | 0.999 | 0.996 | 0.996 | 0.999 | 0.988 | 0.990 | 0.997 |
| HepG2 | 0.997 | 0.999 | 0.999 | 0.995 | 0.997 | 0.999 | 0.987 | 0.993 | 0.996 |
| Jurkat | 0.995 | 0.996 | 0.997 | 0.986 | 0.992 | 0.994 | 0.975 | 0.983 | 0.989 |
| HCT116 | 0.991 | 0.986 | 0.989 | 0.994 | 0.988 | 0.991 | 0.982 | 0.969 | 0.978 |
| HEK293T | 0.972 | 0.983 | 0.974 | 0.882 | 0.980 | 0.923 | 0.795 | 0.956 | 0.858 |

**Table 5: Panel-size sensitivity of the response geometry.**
| (b) Cross-cell concordance at each panel size |  |  |  |
| --- | --- | --- | --- |
| Kendall’s $W$ | 1,000 | 2,000 | 5,000 |
| Alignment $a_{cp}$ | 0.590 | 0.587 | 0.562 |
| Magnitude $m_{cp}$ | 0.524 | 0.533 | 0.546 |
| Coordinate $s_{cp}$ | 0.587 | 0.588 | 0.581 |

**Figure 5:**
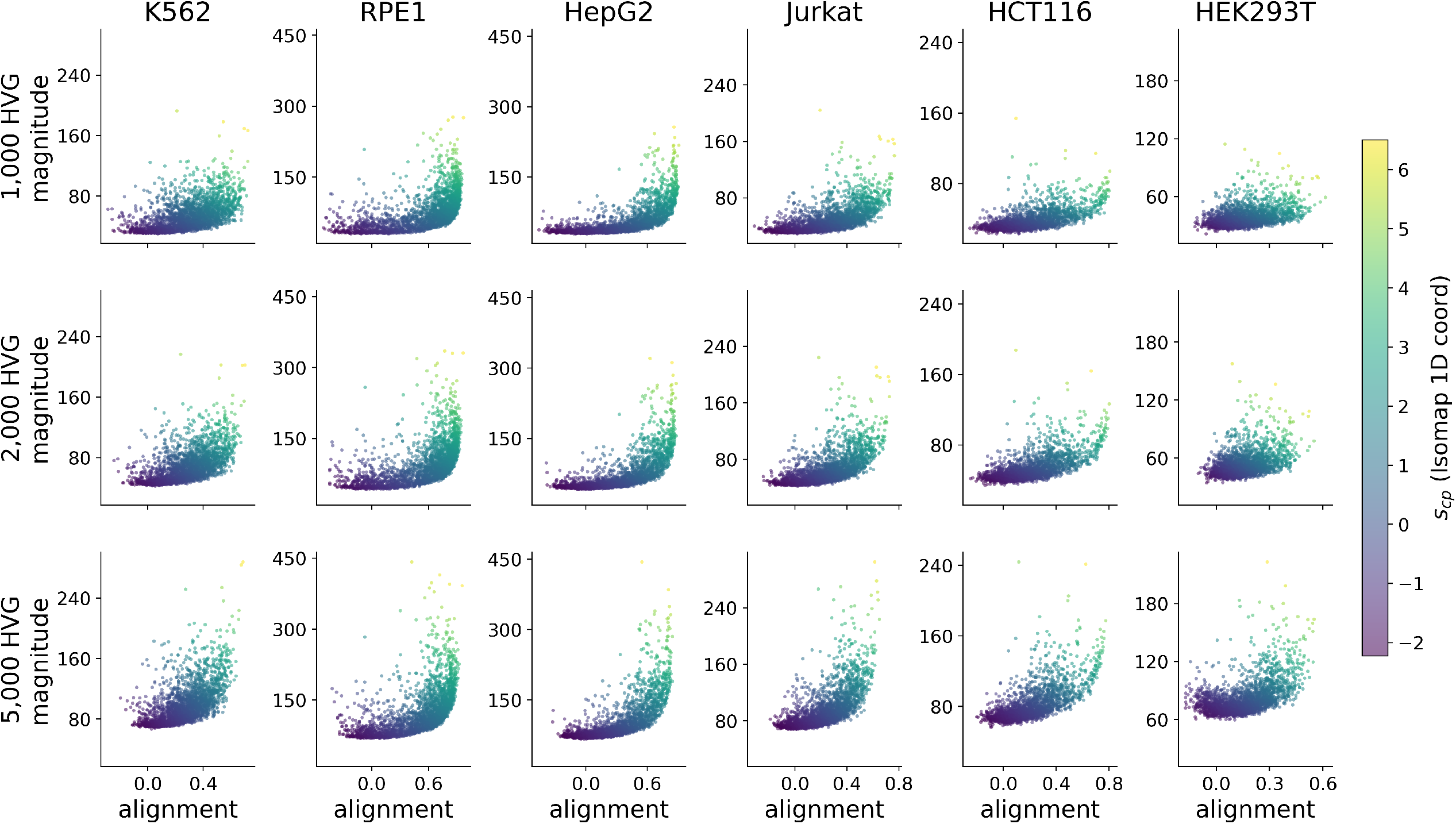
The magnitude–alignment geometry is preserved across readout panel sizes. Each panel plots magnitude *m_cp_* against alignment *a_cp_* for the 2,270 shared perturbations, coloured by the per-cell coordinate *s_cp_*, with rows giving the 1,000-, 2,000-, and 5,000-gene per-cell highly variable panels and columns the six cell lines. Quantitative agreement between panel sizes is reported in Table 5.

### B.3 Gene Ontology enrichment at the two ends and its robustness to the tail cutoff

**Table 6:** Gene Ontology enrichment at the two ends of the response continuum. Perturbations ranked by the cross-cell mean coordinate *s^-^*(*p*); top and bottom 30% tested by gene set enrichment analysis^21^ with Benjamini–Hochberg correction against the genome-wide background.^22^ Terms are Biological Process unless marked *(MF)*. Terms shown are representative specific categories; broad compartment and binding categories such as nucleoplasm, protein binding, and cytosol reach smaller *q* at both ends but are omitted because their size makes them significant against a genome-wide background for almost any gene set. In total 571 terms are significant at the high-alignment end and 293 at the low-alignment end, in which the full list can be found on github. Cutoff robustness results are in Supplementary Table 7.

| High-alignment end (top 30%) |  |  | Low-alignment end (bottom 30%) |  |  |
| --- | --- | --- | --- | --- | --- |
| GO term | hits | $q$ | GO term | hits | $q$ |
| cytoplasmic translation | 71 | $2 \times 10^{-94}$ | regulation of DNA-templated transcription | 56 | $4 \times 10^{-4}$ |
| rRNA processing | 70 | $9 \times 10^{-81}$ | transcription coactivator activity (*MF*) | 30 | $1 \times 10^{-8}$ |
| mRNA splicing via spliceosome | 65 | $2 \times 10^{-49}$ | chromatin remodeling | 27 | $4 \times 10^{-5}$ |
| RNA polymerase II preinitiation complex | 29 | $2 \times 10^{-28}$ | mitochondrial translation | 22 | $4 \times 10^{-14}$ |
| DNA replication | 24 | $4 \times 10^{-17}$ | protein folding | 16 | $6 \times 10^{-4}$ |
| translation initiation factor activity (*MF*) | 18 | $8 \times 10^{-14}$ | ubiquitin-protein transferase activator (*MF*) | 4 | $2 \times 10^{-3}$ |

**Table 7:** The two-end GO enrichment is robust to the tail cutoff. Benjamini–Hochberg *q*-value of each main-text term (Table 6) at three cutoffs of the cross-cell mean coordinate *s^-^*(*p*), using gene set enrichment analysis against the genome-wide background. Every term stays significant at all three cutoffs; several low *s_c_* terms strengthen substantially as the cutoff widens (for example mitochondrial translation, 2 × 10^−3^ to 4 × 10^−14^, and transcription coactivator activity, 2 × 10^−2^ to 1 × 10^−8^), reflecting increased statistical power as the tested set grows rather than a cutoff-specific effect.

| Ontology | GO term | $q$ @ 20% | $q$ @ 25% | $q$ @ 30% |
| --- | --- | --- | --- | --- |
| **High-alignment end** (top tail by cross-cell $\bar{s}(p)$) | | | | |
| BP | cytoplasmic translation | $3 \times 10^{-94}$ | $8 \times 10^{-94}$ | $2 \times 10^{-94}$ |
| BP | rRNA processing | $4 \times 10^{-62}$ | $7 \times 10^{-74}$ | $9 \times 10^{-81}$ |
| BP | RNA polymerase II preinitiation complex | $2 \times 10^{-25}$ | $1 \times 10^{-27}$ | $2 \times 10^{-28}$ |
| BP | mRNA splicing via spliceosome | $7 \times 10^{-24}$ | $1 \times 10^{-41}$ | $2 \times 10^{-49}$ |
| BP | DNA replication | $2 \times 10^{-12}$ | $2 \times 10^{-16}$ | $4 \times 10^{-17}$ |
| MF | translation initiation factor activity | $7 \times 10^{-14}$ | $3 \times 10^{-15}$ | $8 \times 10^{-14}$ |
| **Low-alignment end** (bottom tail by cross-cell $\bar{s}(p)$) | | | | |
| BP | regulation of DNA-templated transcription | $6 \times 10^{-4}$ | $3 \times 10^{-3}$ | $4 \times 10^{-4}$ |
| BP | mitochondrial translation | $2 \times 10^{-3}$ | $5 \times 10^{-5}$ | $4 \times 10^{-14}$ |
| BP | protein folding | $2 \times 10^{-3}$ | $1 \times 10^{-3}$ | $6 \times 10^{-4}$ |
| BP | chromatin remodeling | $7 \times 10^{-3}$ | $2 \times 10^{-3}$ | $4 \times 10^{-5}$ |
| MF | ubiquitin-protein transferase activator | $8 \times 10^{-4}$ | $2 \times 10^{-3}$ | $2 \times 10^{-3}$ |
| MF | transcription coactivator activity | $2 \times 10^{-2}$ | $3 \times 10^{-5}$ | $1 \times 10^{-8}$ |

### B.4 Comparison of gene representations for predicting response position

**Figure 6:**
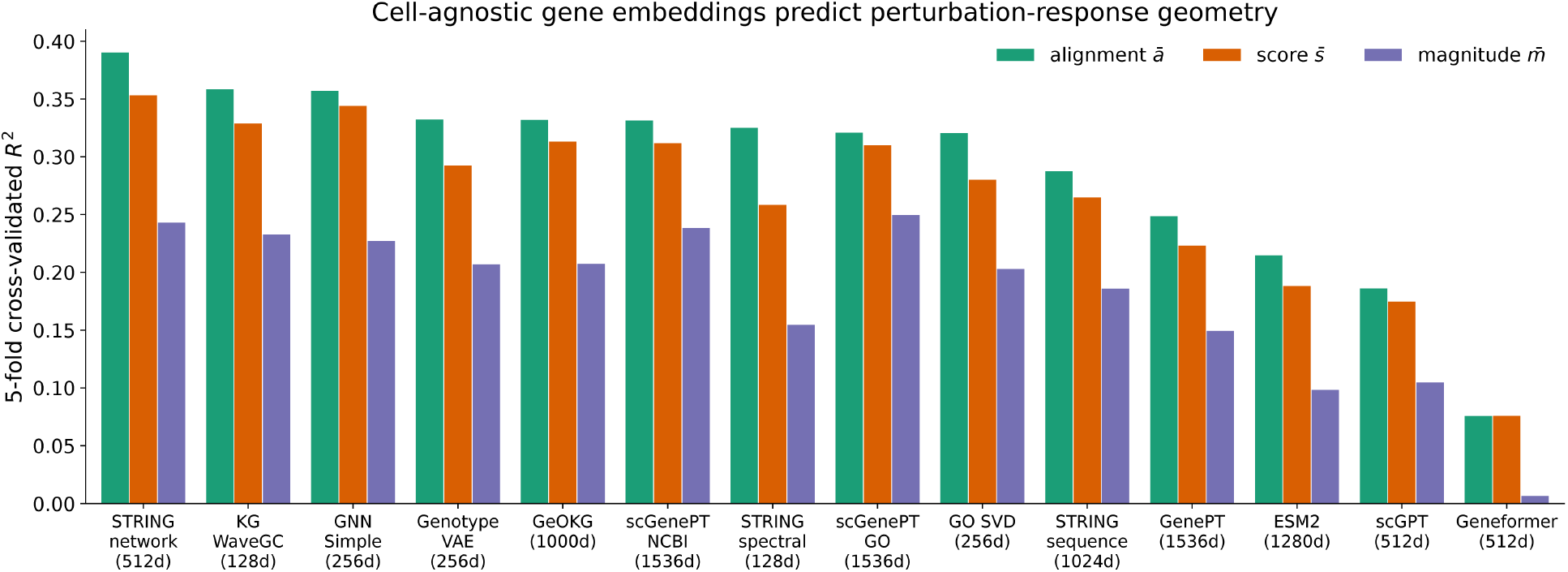
Gene embeddings predict perturbation-response position. Five-fold cross-validated *R*^2^ for predicting cross-cell mean alignment *a^-^*(*p*), magnitude *m^-^* (*p*), and position *s^-^*(*p*) from prior gene representations under identical folds. STRING protein-interaction embeddings are the strongest individual representation and are the interaction space used by COMPASS-N.

### B.5 The fitted coefficient recovers the response geometry

**Table 8:** The fitted shared-response coefficient recovers the geometry defined by the raw coordinate. The analysis of Section 2.6 was repeated with the fitted coefficient *β_p_* in place of the cross-cell coordinate *s^-^*(*p*): the same perturbations ranked, the same top and bottom 30% taken, and the same gene set enrichment analysis with Benjamini–Hochberg correction. The two coordinates use different cell sets, *s^-^*(*p*) averaging over all six lines and *β_p_* over only the four Replogle/Nadig cells that share a readout panel, so their agreement is not built in; they correlate at Pearson 0.88 (Spearman 0.90), and the two tails share 80% and 82% of their perturbations. Every term of Table 6 remains significant when the tails are defined by *β_p_*: eight are more significant, one is unchanged, and three are weaker, DNA replication most noticeably. Across the whole ontology, term-level agreement is high at the high-alignment end (Jaccard 0.77) but much lower at the other end (0.47) despite the similar tail overlap, consistent with a functionally heterogeneous low *s_c_* tail in which small changes in membership alter which of many weakly enriched categories pass the threshold.

| GO term | $q$ from $\bar{s}(p)$ tail | $q$ from $\beta_p$ tail |
| --- | --- | --- |
| **High-alignment end** (top 30%) |  |  |
| cytoplasmic translation | $2 \times 10^{-94}$ | $2 \times 10^{-92}$ |
| rRNA processing | $9 \times 10^{-81}$ | $4 \times 10^{-81}$ |
| mRNA splicing via spliceosome | $2 \times 10^{-49}$ | $4 \times 10^{-61}$ |
| RNA polymerase II preinitiation complex | $2 \times 10^{-28}$ | $1 \times 10^{-28}$ |
| DNA replication | $4 \times 10^{-17}$ | $1 \times 10^{-12}$ |
| translation initiation factor activity | $8 \times 10^{-14}$ | $3 \times 10^{-15}$ |
| **Low-alignment end** (bottom 30%) |  |  |
| regulation of DNA-templated transcription | $4 \times 10^{-4}$ | $1 \times 10^{-4}$ |
| transcription coactivator activity | $1 \times 10^{-8}$ | $2 \times 10^{-7}$ |
| chromatin remodeling | $4 \times 10^{-5}$ | $2 \times 10^{-8}$ |
| mitochondrial translation | $4 \times 10^{-14}$ | $5 \times 10^{-23}$ |
| protein folding | $6 \times 10^{-4}$ | $6 \times 10^{-4}$ |
| ubiquitin-protein transferase activator | $2 \times 10^{-3}$ | $1 \times 10^{-3}$ |
| Across the whole ontology | High-alignment | Low-alignment |
| Significant terms from $\bar{s}(p)$ | 571 | 293 |
| Significant terms from $\beta_p$ | 569 | 236 |
| Shared | 496 | 169 |
| Term-level Jaccard | 0.77 | 0.47 |

### B.6 Transfer of the decomposition with a single source cell line

**Table 9:**
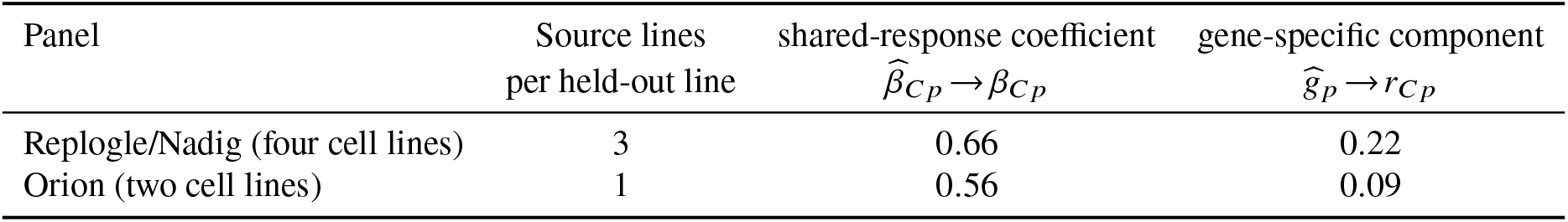
Leave-one-cell-out transfer degrades when only one source cell line is available. Each held-out cell line receives a coefficient and a gene-specific component estimated from the remaining lines in its study, and both columns report agreement with the held-out cell line’s own value: Pearson correlation across the 2,270 shared perturbations for the coefficient, and the mean per-perturbation Pearson correlation across genes for the gene-specific response, as in Table 2. The Replogle/Nadig row averages the four held-out cell lines in Table 2. The Orion row averages its two cell lines; these values are identical by symmetry because, with only two lines, each source-cell average is the single other cell. Having only one donor cell line costs the gene-specific component far more than the coefficient.

### B.7 Blend-weight landscape for the cross-context predictor

**Figure 7:**
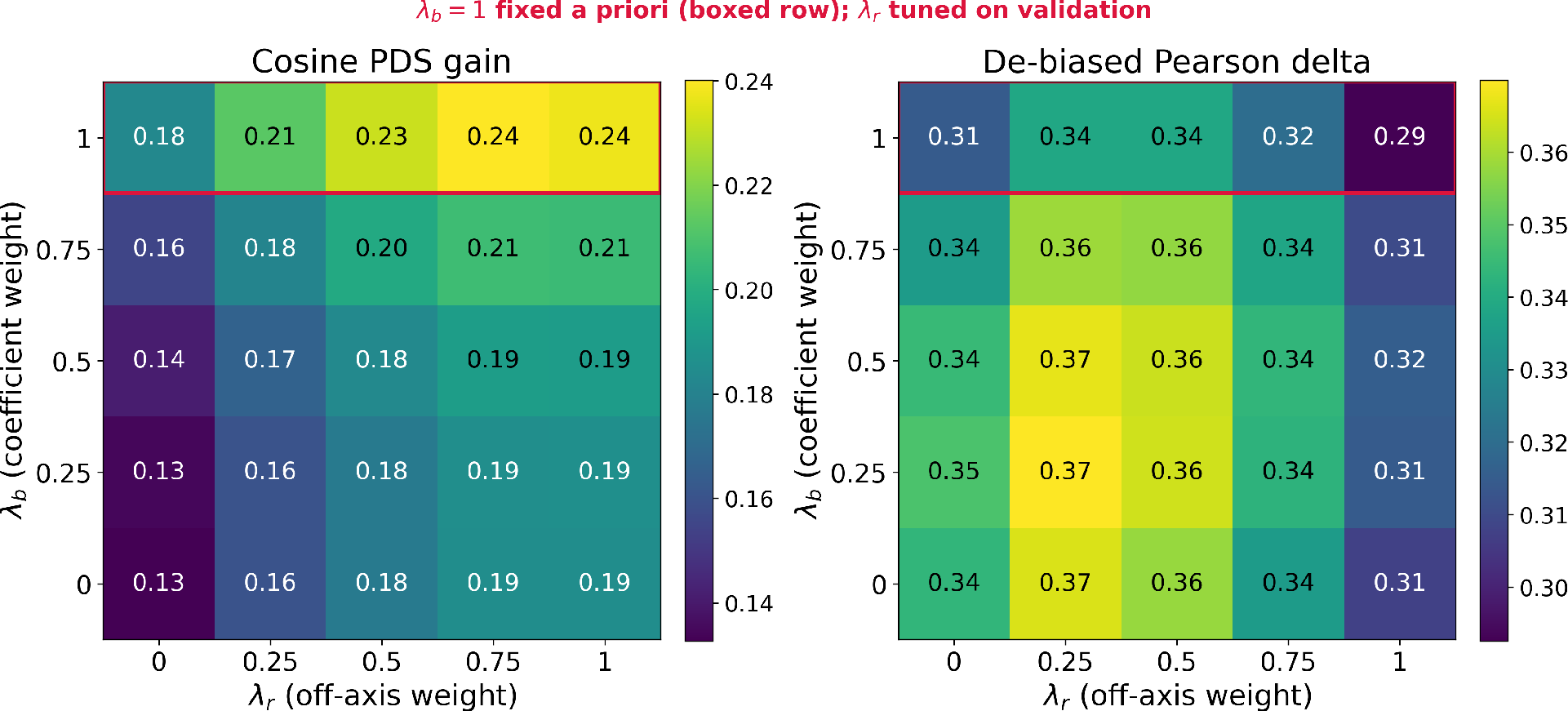
Blend-weight landscape. (*λ_b_*, *λ_r_*) **for COMPASS-H, pooled over the six cell lines.** *λ* = 1 uses the cross-context estimate and *λ* = 0 the in-context estimate. Cosine PDS gain increases with *λ_b_* and peaks at *λ_b_* = 1, whereas de-biased Pearson delta peaks near *λ_b_* = 0.25 because the in-context coefficient is closer in magnitude. We fix *λ_b_* = 1 rather than tuning it: the cross-context estimate of the shared-response coefficient is measured on the same perturbation in other cell lines, so it preserves that perturbation’s shared-response coefficient, and it is this ordering that discrimination depends on. Trading the Pearson-optimal value for *λ_b_* = 1 costs about 0.03 in correlation and gains about 0.05 in PDS. The preferred *λ_r_* differs across metrics, so it is selected on the validation partition using a finer grid than shown.

### B.8 Comparison of the COMPASS suite as in-context training perturbations increase

**Figure 8:**
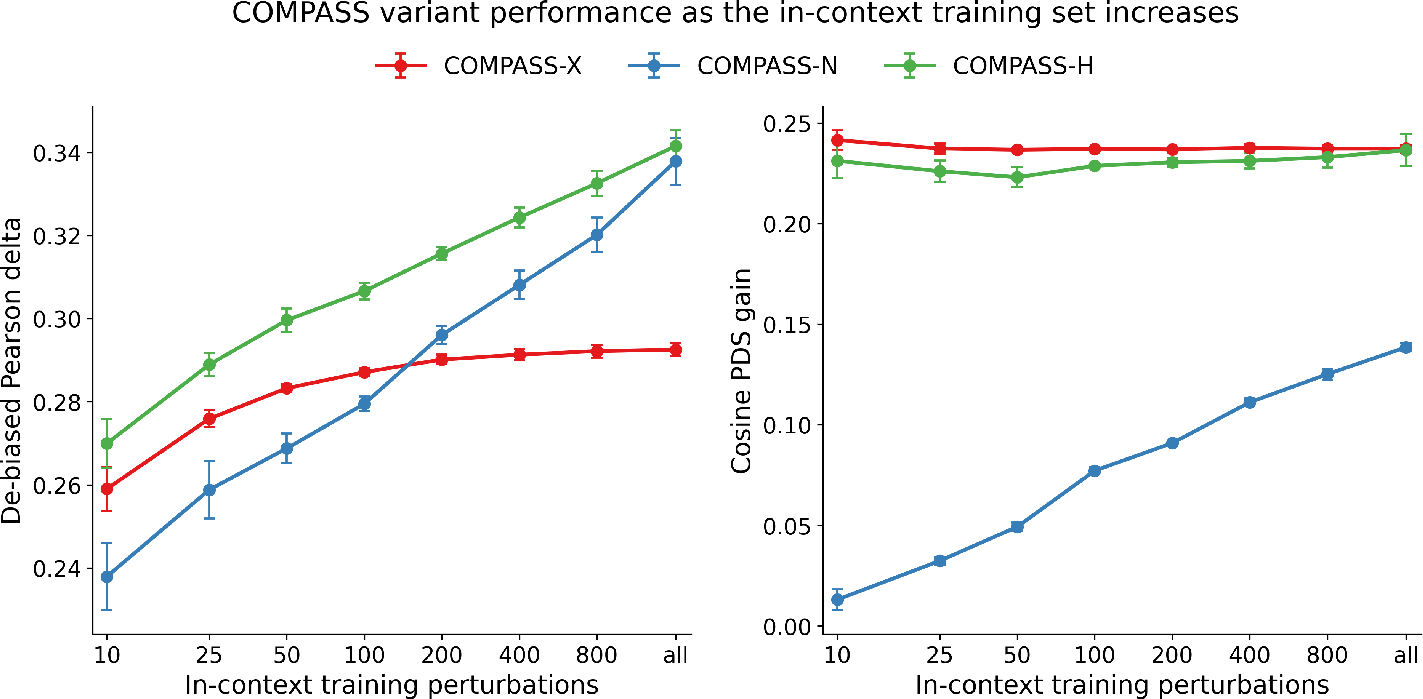
Low-data sweep. De-biased Pearson delta (left) and cosine PDS gain (right) as the number of in-context training perturbations in the query cell line varies over {10, 25, 50, 100, 200, 400, 800, all} (log scale; “all” ≈ 1,620 training perturbations). Averaged over six cell lines and 15 draws per training size (five random train/test partitions, each subsampled three times), except at “all”, where the training set is fixed and only the five partitions vary; error bars are the standard error across cell lines after removing each cell line’s own mean across training sizes.

### B.9 gene-specific programs cluster into coherent pathways

**Table 10:**
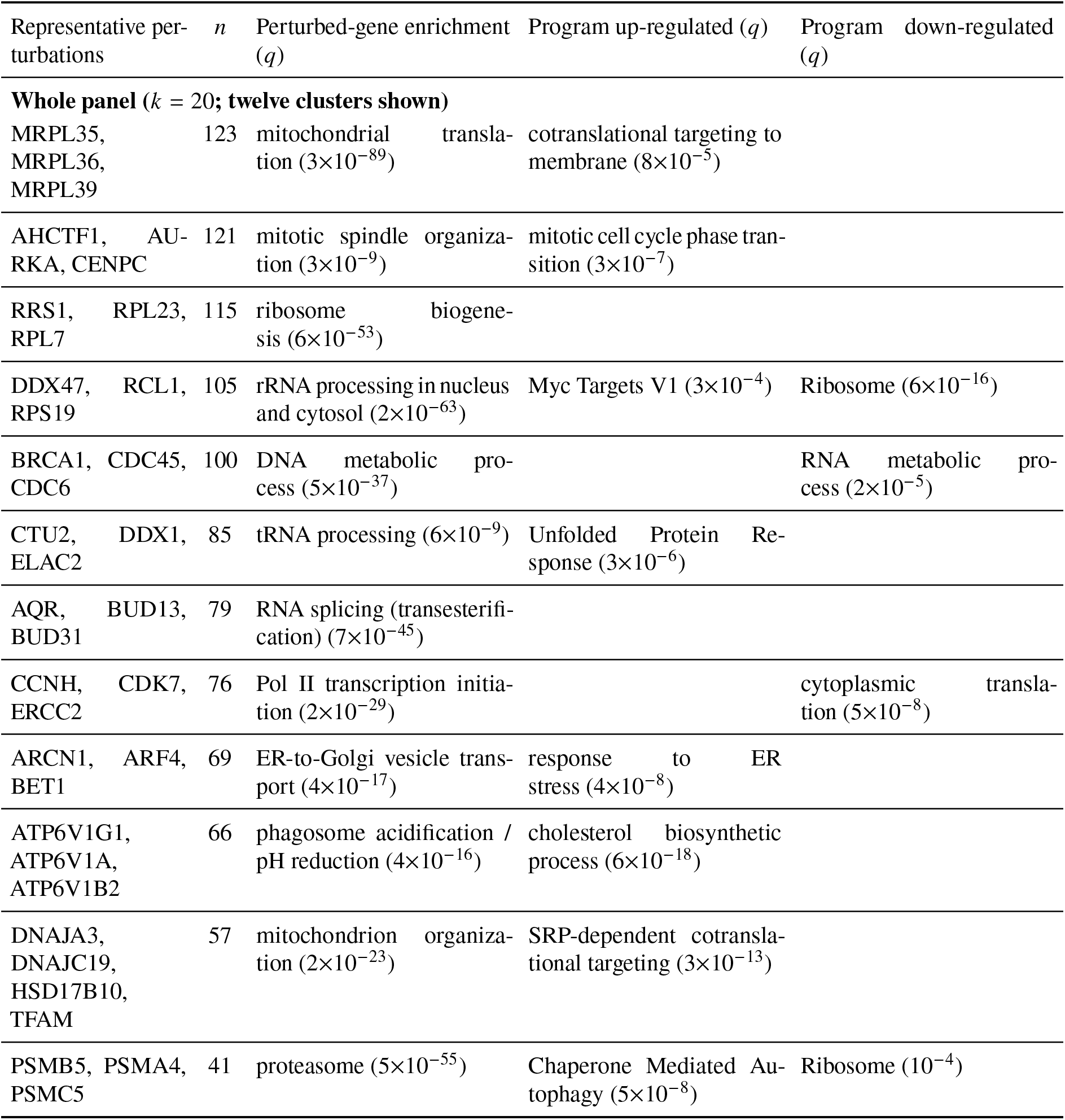
Gene-specific programs(*g_p_*) cluster into coherent pathways (whole panel). The 2,270 perturbations were clustered by the direction of their cross-cell-averaged gene-specific component *g_p_* (*k*-means on L2-normalized *g_p_*, with the number of clusters *k* = 20 fixed a priori rather than chosen by a stability criterion). Each cluster is characterized on two sides: enrichment of its member perturbed genes against the 2,270-perturbation anchor, and enrichment of the fifteen most up- and down-regulated genes of the cluster-mean *g_p_* program against the 2,000-gene readout panel (a fixed gene count, unrelated to *k*). Both use gene set enrichment analysis with Benjamini–Hochberg correction against the Enrichr GO Biological Process 2021, KEGG 2021, Reactome 2022, and MSigDB Hallmark 2020 libraries. Each entry is a representative significant term with its *q*, chosen for interpretability rather than strictly by rank; representative perturbations are cluster members. Twelve of the twenty clusters are shown. Many clusters are coherent on both sides.

**Table 11:**
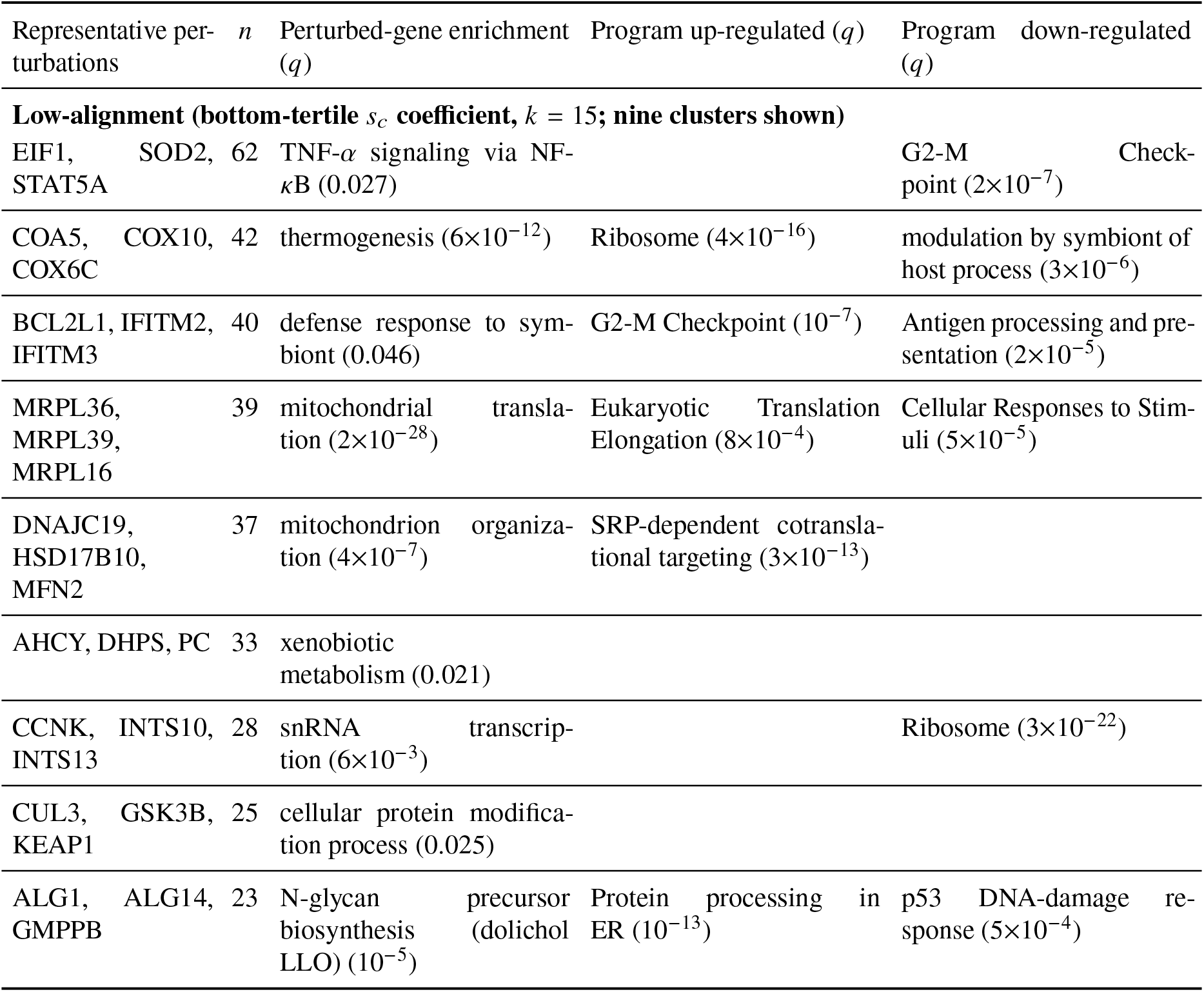
Gene-specific programs cluster into coherent pathways (low-alignment). The analysis follows Table 10 but is restricted to bottom-tertile *s_c_*-coefficient, clustered into *k* = 15 groups, which engage the shared response weakly. The nine clusters whose members reach a significant perturbed-gene enrichment (*q* < 0.05) are shown; the remaining six do not, so even at this end of the axis a majority but not all of the clusters share a dominant perturbed-gene family and a coherent *g_p_* program. As in Table 12, representative perturbations are three of the cluster members that drive its perturbed-gene enrichment.

**Table 12:**
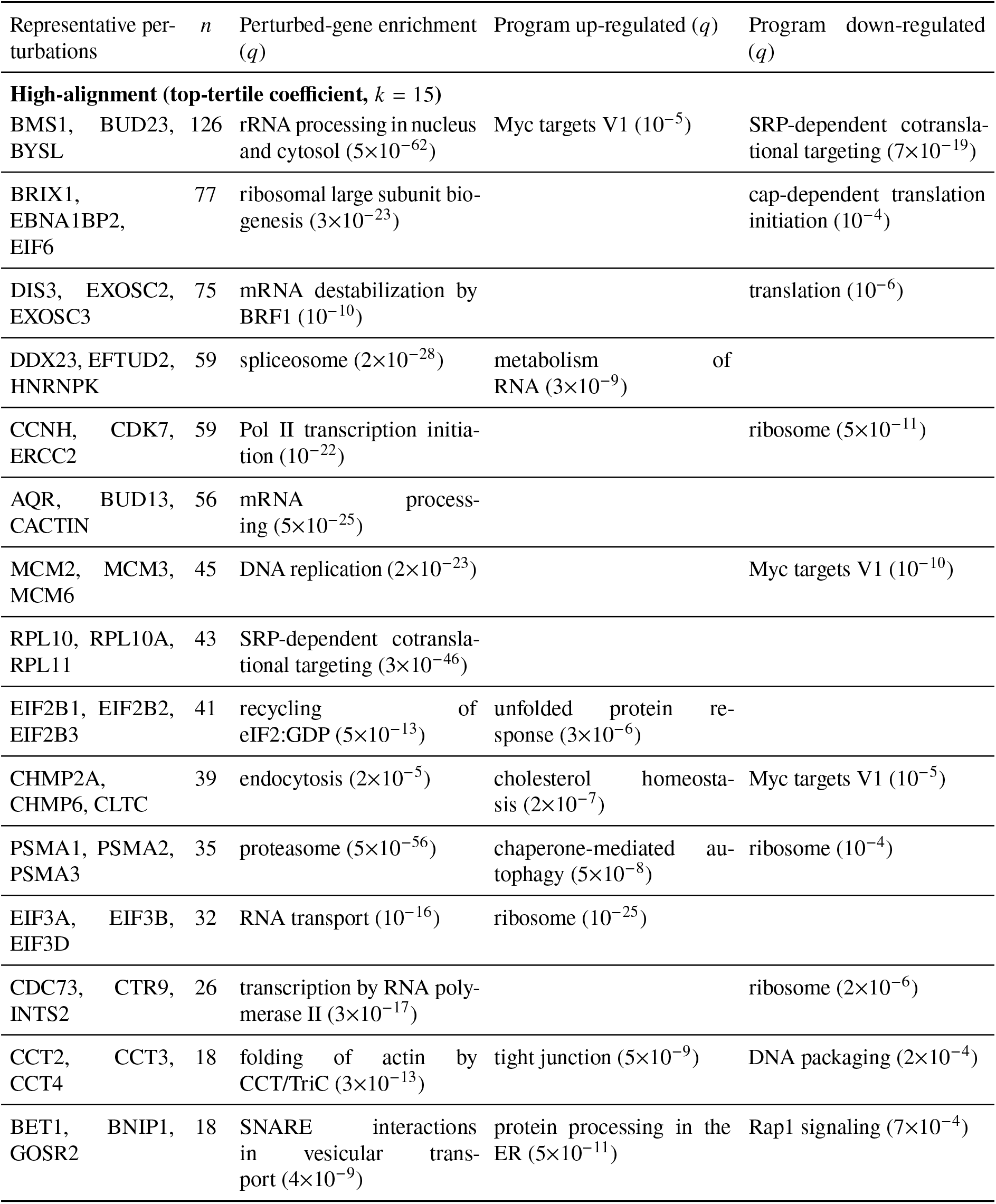
Gene-specific programs cluster into coherent pathways. As Table 10, restricted to top-tertile-coefficient of *s_c_*. Both tertiles hold 749 perturbations and are clustered into the same *k* = 15 groups, so the two ends are directly comparable: every high-*s_c_* cluster reaches a significant perturbed-gene enrichment, against nine of fifteen at the low-*s_c_* end (Table 11). Representative perturbations are three cluster members driving the enrichment; program columns are blank where no term reaches *q* < 10^−3^.

### B.10 The estimated shared-response coefficient is conserved across cell lines

**Table 13:** Cross-cell conservation of the shared-response direction coefficient. Each cell is fit independently, and rows report correlation across perturbations between the resulting *β_cp_*.

| Cell-line pair | Pearson correlation of $\beta_{cp}$ |
| --- | --- |
| *Replogle/Nadig common 2,000-gene panel* |  |
| K562 / RPE1 | +0.50 |
| K562 / HepG2 | +0.54 |
| K562 / Jurkat | +0.56 |
| RPE1 / HepG2 | +0.66 |
| RPE1 / Jurkat | +0.54 |
| HepG2 / Jurkat | +0.53 |
| Mean (six pairs) | +0.55 |
| *Orion common 2,000-gene panel* |  |
| HCT116 / HEK293T | +0.56 |

